# Salivary microRNA Profiling of Long COVID Subjects Reveals Host-Encoded Regulators of Inflammation and Viral Persistence

**DOI:** 10.64898/2026.06.07.730729

**Authors:** Kristelle J. Capistrano, Raza Ali Naqvi, Sarah Elshourbagy, Jake Class, Justin M. Richner, Sodabeh Etminan, Joel L. Schwartz, Wei Li, Christine D. Wu, Afsar R. Naqvi

**Author notes:** Corresponding Author: Afsar Naqvi, Ph.D., Professor, Department of Periodontics College of Dentistry, University of Illinois Chicago, 801 S. Paulina, Room 561B, Chicago, IL 60612.

## Abstract

Periodontal disease and COVID-19 are linked by convergent immunoinflammatory pathways, yet the molecular basis of their interaction remains poorly defined. Here, we present a comprehensive salivary microRNA profile from individuals with prior SARS-CoV-2 infection, sampled approximately 3–6 months after diagnosis and meeting criteria for long COVID, providing new insight into the post-viral oral microenvironment. Salivary miRNA sequencing revealed widespread repression in patients with PD, consistent with persistent immune dysregulation. Relative to COVID-19-negative/PD-negative controls, thirty-two miRNAs were differentially expressed in COVID-19-positive/PD-positive individuals, all significantly downregulated. A similar signature was observed in a post-vaccination cohort for the selected dysregulated miRNAs. Integrative pathway analyses identified these miRNAs as regulators of core inflammatory circuits, including Ras, MAPK, and NFκB signaling, converging on IL-1β- and TNF-centered networks relevant to both PD and COVID-19. Mechanistically, restoration of three downregulated miRNAs, miR- miR-30e-3p 106-3p-3p, and miR-652-3p attenuated NFκB activation and cytokine release in TLR-stimulated human oral keratinocytes, while their functional suppression using inhibitors potentiates inflammation. These miRNAs were also predicted to target SARS-CoV-2 spike and nucleocapsid transcripts, an interaction validated by dual-luciferase reporter assays. Their overexpression further reduced spike and nucleocapsid expression in Beta- and Omicron-infected epithelial cells, as measured by flow cytometry and RT-qPCR confirming host miRNAs as potent endogenous SARS-CoV-2 restriction factor. Together, these findings identify salivary host miRNAs as mechanistic regulators of oral inflammatory tone and viral persistence, establishing a molecular link between periodontal inflammation and post-COVID oral pathology.

## INTRODUCTION

Despite the World Health Organization declaring the end of the COVID-19 pandemic in 2023, the long-term consequences of disease continue to pose significant health challenges, encompassing over 200 reported symptoms that include persistent fatigue, shortness of breath, neurocognitive impairment, dyspnea, and myocardial inflammation, necessitating continued clinical monitoring and coordinated long-term care.^1^ Individuals who develop or continue to experience these symptoms at least three months post the initial SARS-CoV-2 infection suffer from Long COVID (LC). Current estimates suggest that 1 out of 10 COVID-19 patients exhibit LC signs and symptoms.^2,3^ Notably, SARS-CoV-2 can persist in the oral cavity, where the virus has been shown to disrupt mucosal immunity and contribute to oral pathologies as LC.^4–9^ Among these pathologies, we focus on periodontal disease (PD), a microbial biofilm-driven condition that damages the tooth-supporting structures, including the gingiva, periodontal ligament (PDL), cementum, and alveolar bone. High prevalence of PD (∼42% of US population)^10^ and its emerging association with COVID-19, driven by the identification of the periodontium as a potential reservoir for SARS-CoV-2^11,12^ and shared immune dysregulation makes this bidirectional relationship biologically of broad interest. These associations underscore the need to explore the underlying molecular mechanisms linking PD and LC.

The periodontium plays a dual role in SARS-CoV-2 infection, serving not only as a portal of entry but also as a site that supports viral attachment, replication, and shedding.^12^ Cells expressing the primary SARS-CoV-2 entry receptor angiotensin-converting enzyme 2 (ACE2), along with the host proteases furin and transmembrane serine protease 2 (TMPRSS2) that activate the spike protein, abound in the periodontium, particularly in patients with periodontitis.^12–15^ Of note, RT-PCR analysis of postmortem biopsies from COVID-19 fatalities revealed SARS-CoV-2 RNA in periodontal tissues.^13^ These findings implicate PD in SARS-CoV-2 entry and persistence. Virulence factors expressed by dysbiotic periodontopathic bacteria in PD may further contribute to viral adhesion and entry conditioning periodontal tissues for long term viral persistence.^5,16^

A key hallmark shared by PD and COVID-19, and central to their bidirectional relationship, is immune dysregulation, characterized by the chronic overproduction of proinflammatory cytokines, chemokines, and other mediators, particularly in subjects with moderate to severe PD.^5,17^ Both conditions activate overlapping inflammatory pathways such as NFκB, NLRP3/IL-1β, IL-6, and MAPK, which, when sustained, can drive tissue destruction and, in severe cases, organ damage.^18^ Chronic periodontal inflammation can also extend beyond local tissues and elevate systemic inflammatory burden.^17,19^ Conversely, acute phase of COVID-19 associates with heightened release of inflammatory cytokines (i.e., IL-1β, IL-6, and TNF□), potentially increasing periodontal inflammation.^20,21^ These findings suggest that PD may play a compounding role in COVID-19 pathogenesis and highlight the potential of SARS-CoV-2 to exert lasting effects on oral health.^9,22^ Given the clinical and molecular associations between periodontal disease (PD) and COVID-19, it is imperative to develop treatments that target their interplay. This requires a deeper understanding of the mechanisms driving additive immune dysregulation. One such mechanism may involve host microRNAs (miRNAs), which remain poorly understood within the context of PD and COVID-19. This knowledge gap is significant, as miRNAs are potent, endogenous post-transcriptional regulators of immune responses. These small, single-stranded, non-coding RNAs (19–24 nts long) function by binding complementary sequences on target mRNAs, resulting in transcript degradation or translational repression. Studies have shown that miRNAs are differentially expressed in both PD ^23–25^, and COVID-19 ^26,27^, and they may play a key role in modulating the inflammatory environment in both conditions. Beyond regulating host immunity, miRNAs reportedly suppress RNA virus replication.^28–30^ Our prior work, along with studies by others, have identified putative miRNA binding sites within the SARS-CoV-2 genome, particularly in the receptor binding domain (RBD) of S protein, thereby limiting viral replication and release.^31–33^ The SARS-CoV-2 nucleocapsid (N) protein, which is essential for viral replication and immune evasion, also presents a promising miRNA target. While several studies have explored the role of miRNAs in either COVID-19 or PD independently, to our knowledge, no study has examined salivary miRNA profiles within the integrated context of both conditions. More importantly, the mechanisms underlying miRNA regulation of SARS-CoV-2 oral tropism remains poorly explored. Our study addresses these gaps by investigating LC and PD associated miRNAs that play crucial role in SARS-CoV-2 tropism and oral inflammation. We identified miRNAs dysregulation in SARS-CoV-2+/PD+ (LC), which impaired antiviral and inflammatory pathways in the oral cavity.

## MATERIALS and METHODS

### Patient Data

This study was approved by the Institutional Review Board of the University of Illinois at Chicago (UIC IRB#2016-0696) and the Miles Square Health Center Research Council (approved December 2020). All procedures adhered to the principles of the Declaration of Helsinki and HIPAA regulations. Ethical oversight encompassed both the retrospective review of clinical data and the prospective collection of biological specimens, including saliva and gingival crevicular fluid.

#### Inclusion and Exclusion Criteria

Inclusion criteria included adults aged 21–70 years who had abstained from oral activity (e.g., eating, drinking, rinsing, or tooth brushing) for ≥1 hour prior to sampling and had no antibiotic use within one week. To the best of our ability, we aimed to include an equal number of males and females (sex assigned at birth). Exclusion criteria included individuals with an active SARS-CoV-2 infection.

### Sample collection and processing

Full-mouth periodontal examinations were carried out by calibrated examiners at either the University of Illinois at Chicago College of Dentistry (UIC-COD) or Mile Square Health Center from December 2021 (Pre-vaccination) to March 2024 (Post-vaccination). Each tooth was as previously described.^34^

COVID-19 history was self-reported. Stimulated saliva was collected by having participants chew sterile paraffin wax for 5 minutes and expectorate into a sterile 50 mL conical tube (n=10/group; SARS-CoV-2+/PD+ (LC), SARS-CoV-2+/PD-, SARS-CoV-2-/PD-). The volume and physical characteristics of each sample were documented, with salivary flow rates below 5 mL per 5 minutes indicating reduced secretion (xerostomia).

Sample consistency was also noted, ranging from watery to mucoid or stringy-viscous. Saliva samples were then stored in 1:1 volume of QIAzol reagent (Qiagen, Hilden, Germany) and stored at –80°C until further processing.

### miRNA Sequencing

Total RNAs, including miRNAs, were extracted from 500 µL of saliva using commercial miRNeasy Qiagen RNA isolation following the manufacturer’s protocols (Qiagen, Hilden, Germany). RNA quality was verified using an Agilent Bioanalyzer 2100 system, and only samples with RNA Integrity Number (RIN) greater than 7.0 were included for downstream analysis. Small RNA library construction and sequencing were performed by LC Sciences (Houston, TX, USA) as part of their miRNA sequencing service. Bioinformatic analysis was performed using the ACGT101-miR pipeline (LC Sciences), which includes standard procedures for adapter trimming, filtering, and alignment to miRBase v22.0.

### Computational Prediction of SARS-CoV-2 Target Sequences

#### RNAHybrid 2.0, Multiple Sequence Alignment Analysis (MUSCLE), and NIH Nucleotide

We obtained SARS-CoV-2 sequence information from GISAID: B.1.351 (EPI_ISL_745146) and B.1.1.529 (EPI_ISL_6841980). Sequences of significantly downregulated and known mature miRNAs in SARS-CoV-2+/PD+ (LC) versus SARS-CoV-2-/PD- (**Table 1**) were obtained from miRbase database version 22.1. Predicted binding sites of these miRNAs on B.1.351 and B.1.1.529 spike and nucleocapsid sequences were computationally determined through RNAhybrid 2.2. Binding parameters for miRNA:spike predictions were set as the following: 4 hits per target, −25 kcal/mol energy threshold, helix constraint = 2–8, max bulge loop length = 2, and p value = nothing. miRNA:nucleocapsid binding parameters were set as: 4 hits per target, −10 kcal/mol energy threshold, helix constraint = 2–8, max bulge loop length = 2, p value = nothing.

MUSCLE alignment was used to identify conserved spike transcript binding regions ^31^, followed by Nucleotide validation (https://www.ncbi.nlm.nih.gov/nuccore).

### Subcloning SARS-CoV-2 Spike and Nucleocapsid Transcripts into Reporter Plasmid

#### Design and Synthesis of Custom Nucleocapsid Plasmid

We designed custom vector systems to express SARS-CoV-2 nucleocapsid gene sequences from B.1.351 (GISAID: EPI_ISL_745146) and B.1.1.529 (GISAID: EPI_6841980). Each sequence was synthesized with flanking XhoI and NotI restriction sites and assembled into GeneArt^TM^ pMA-RQ cloning vector (ThermoFisher Scientific, Wilmington, MA) using standard molecular cloning techniques. The sequence of these custom plasmids can be found in **Fig.S3**.

#### PCR Amplification of SARS-CoV-2 Spike

Plasmids encoding the full-length SARS-CoV-2 spike (Invivogen pUNO1-SpikeV07: B.1.351 (GISAID: EPI_ISL_745146) and Invivogen pUNO1-SpikeV11: B.1.1.529 (GISAID: EPI_ISL_6841980)) and nucleocapsid transcripts were PCR-amplified using high-fidelity DNA polymerase (Q5® High-Fidelity DNA Polymerase, New England Biolabs, MA). Primers were designed to incorporate XhoI and NotI restriction enzyme sites at the 5’ and 3’ ends, respectively (**Fig.S3**). PCR products were then purified using QIAquick PCR Purification Kit (Qiagen, Hilden, Germany), followed by agarose gel electrophoresis to confirm size specificity (B.1.351 spike: 3757 bp and B.1.1.529 spike: 3755 bp; B.1.351 nucleocapsid: 1251 bp and B.1.1.529 nucleocapsid: 1390 bp).

#### Restriction Digestion, Ligation, and Transformation

Purified PCR products and psiCHECK-2 reporter plasmid (6273 bp) were digested according to the manufacturer’s instructions (FastDigest XhoI and NotI, ThermoFisher Scientific, Wilmington, MA) at 37°C for 30 minutes. The digested insert and vector were purified using QIAquick PCR Purification Kit (Qiagen, Hilden, Germany). Sticky-end ligation was performed using T4 DNA ligase (NEB, Ipswich, MA) at a 4:1 molar ratio of insert to vector. Sanger sequencing further confirmed positive clones. These plasmids (psiCHECK-2-SARS-CoV-2-S and psiCHECK2-SARS-CoV-2-N) were then transformed into NEB 10-beta chemically competent *E. coli* following the manufacturer’s protocol (NEB, Ipswich, MA).

### Cell Transfection and Dual Luciferase Assay

To validate direct binding of our candidate miRNAs to SARS-CoV-2 spike (miR-106b-3p, miR-106B-5p, miR-425-5p, miR-652-3p, and miR-338-3p (negative control)) or nucleocapsid transcripts (miR-30e-3p, miR-378d, miR-223-5p, miR-223-3p, miR-652-3p, and miR-425-5p (negative control)), we first co-transfected mirVana^TM^ miRNA mimics (Invitrogen, Waltham, MA) at 1, 2.5, or 10 pmol and psiCHECK-2-SARS-CoV2-S or psiCHECK-2-SARS-CoV-2-N (50 ng) into 293T Human Epithelial Keratinocytes according to the manufacturer’s instructions (Lipofectamine^TM^ 2.0 Transfection Reagent, Invitrogen, Waltham, MA). Thirty-six hours post miRNA mimic and reporter vector co-transfection, the cell lysates underwent the Dual-Glo Luciferase assay system (ProMega, Madison, WI) according to the user manual. Relative Luciferase activities were normalized to control mimic luciferase activities and determined by calculating the ratio of Firefly luciferase activities over Renilla luciferase activities. Experiments were repeated four times as quadruplets on a white Greiner CELLSTAR 96-well Plate (x). Readings were obtained on a luciferase plate reader (Biotek, Winooski, VT, USA).

### KEGG and IPA

KEGG pathway analysis on differentially expressed miRNAs was performed by LC Sciences as part of its miRNA sequencing service. Subsequent bioinformatic analyses were performed using Ingenuity Pathway Analysis (IPA) software (© QIAGEN 2013–2024). **Fig.4C-D** were generated using the IPA Path Designer tool, in which biological reactions between molecules are depicted as edges (lines).

### Transfection Using miRNA Mimics and Overexpression Plasmids

We performed transient transfections of (**a**) mirVana^TM^ miRNA mimics (50nM)-miR-30e-3p, miR-106b-3p, miR-652-3p, and negative control #1 or (**b**) miRNA overexpression plasmids (250 ng)- miR-30e, miR-106b, and miR-652 (OriGene, Rockville, MD)-into SARS-CoV-2-permissive ACE2-A549 epithelial cells using Lipofectamine 2000 (Invitrogen, Waltham, MA) for 36 hours according to the manufacturer’s instructions.

### Viral Infection of Transfected Cells

Cells transfected with either miRNA mimics or overexpression plasmids were challenged with 0.5 MOI of SARS-CoV-2 B.1.351 (Source: BEI (NR-55282)) or B.1.1.529 (Source: BEI: NR-56461)) for 24 or 48 hours. After each timepoint, cells for RNA extraction were stored in 700 μL of QIAzol lysis reagent (Qiagen, Hilden, Germany) at −80°C, whereas cells for flow cytometry analysis were fixed in 2% PFA and stored in 4°C until further analysis.

### Quantification of SARS-CoV-2 Spike Protein and Nucleocapsid mRNA Expression

#### RT-qPCR

After viral infection, the supernatant was discarded and SARS-CoV-2-infected and miRNA-transfected cells were washed 3 times in 1X PBS. Cells intended for RT-qPCR were stored in 700 uL of QIAzol reagent (Qiagen, Hilden, Germany) and stored in – 80°C until further processing. Total RNA of SARS-CoV-2-infected cells were extracted using commercial miRNeasy Qiagen RNA isolation following the manufacturer’s protocols (Qiagen, Hilden, Germany). Real-time RT-qPCR was performed using the TaqMan® One-Step RNA-to-Ct Kit (ThermoFisher Scientific, Wilmington, MA) as previously described.^35^

#### Flow Cytometry

Cells intended for flow cytometry analysis were washed 3 times in 1X PBS, resuspended in 4% PFA, and stored in 4°C until further processing. Flow cytometry was performed on approximately 300,000 cells to evaluate SARS-CoV-2 spike (S) protein expression on the surface and intracellularly in ACE2-A549 cells using anti-SARS-CoV-2 Spike antibody (clone GT263, Invitrogen). Data from 100,000 events were acquired using a Cytek flow cytometer and analyzed with FlowJo_v10.9.0 software.

### microRNA quantification

For miRNA quantification, miRNA-specific primers and the reverse transcription kit were obtained from Qiagen. Reverse transcription was performed using 100 ng of total RNA according to the manufacturer’s instructions. qPCR reactions were carried out using miRNA-specific primers and a universal primer in the reaction mixture. RNU6B served as the endogenous control, and relative fold changes were determined from replicate Ct values using the ΔΔCt method.

### TLR Stimulation and Phospho Flow

To measure phosphorylation of downstream TLR signaling transcription factors, HOKs were fixed 30 min after TLR stimulation at 4 °C. Cells were washed twice with RPMI supplemented with 1% BSA and fixed in Cytofix Buffer (BD Biosciences). Following permeabilization, cells were washed twice more with RPMI-1% BSA and stained with phospho-specific antibodies. Phosphorylation was assessed using antibodies against NFκB (BV421-conjugated anti-NFκB p65, clone K10-895.12.50, BD Biosciences).

### Cytokine Analysis

Supernatants were collected from miRNA or control mimic or inhibitor transfected HOK challenged with TLR4 agonist at 4 and 24 h. Multiplex analysis of four different cytokines (IL-1β, IL-6, IL-8 and TNF-α) was performed using Milliplex (Millipore, Billerica, MA, USA). Data was collected on Bio-Plex flow cytometer (Bio-Rad, Hercules, CA, USA) for analysis.

### Statistical Analysis

Statistical tests were performed using Prism 9 software (GraphPad). Comparisons between two groups were conducted using Student’s t-test on normalized expression values. For experiments involving more than two groups, one-way ANOVA was used to assess statistical significance. For RT-qPCR-based genome quantification, one-way ANOVA with Kruskal-Wallis test was applied.

## RESULTS

### Global Downregulation of Salivary miRNA Expression in Patients with Long COVID

To investigate potential alterations in the oral molecular microenvironment associated with periodontitis (PD) and prior SARS-CoV-2 infection, we performed miRNA-seq on salivary samples from three patient cohorts stratified by PD status and history of SARS-CoV-2 infection (∼3-6 months after initial infection): (1) SARS-CoV-2-/PD-, (2) SARS-CoV-2+/PD-, and (3) SARS-CoV-2-/PD+ (LC). Patients with differential expression analysis revealed distinct patterns of miRNA upregulation and downregulation across the cohorts (**Fig.1A**). In SARS-CoV-2-/PD+ patients compared to healthy (SARS-CoV-2-/PD) controls, we observed 51 upregulated and 49 downregulated miRNAs (p<0.05; **Fig.1B**). In contrast, the SARS-CoV-2+/PD+ (LC) (LC) cohort exhibited a predominant downregulation of miRNAs compared to the other groups. Specifically, when compared to SARS-CoV-2-/PD+ patients, 25 miRNAs were significantly upregulated and 72 were downregulated in the SARS-CoV-2+/PD+ (LC) (LC) group (p < 0.05; **Fig.1B**). Similarly, comparison with healthy controls revealed 31 upregulated and 65 downregulated miRNAs in SARS-CoV-2+/PD+ (LC) patients (p < 0.05; **Fig.1A-B**). This global downregulation was especially pronounced when comparing SARS_CoV-2+/PD+ (LC) to healthy controls at p < 0.01, where no known miRNAs were upregulated; only novel miRNAs were identified as upregulated (**Fig.1C**, **Table 1**). Here, the most downregulated known miRNAs were let-7f-2-3p (fold change=0.04), miR-338-3p (fold change=0.04), miR-378d (fold change=0.05), miR-194-5p (fold change=0.08), and miR-2355-5p (fold change=0.08) (**Table 1**). This pattern was not observed in other pairwise comparisons, such as SARS-CoV-2-/PD+ vs. SARS-CoV-2+/PD+ (LC) or SARS-CoV-2-/PD+ vs. SARS-CoV-2-/PD- (p < 0.01; **Fig.1C**). Collectively, these findings highlight a broad suppression of salivary miRNA expression in individuals with both PD and prior SARS-CoV-2 infection.

**Fig. 1.**
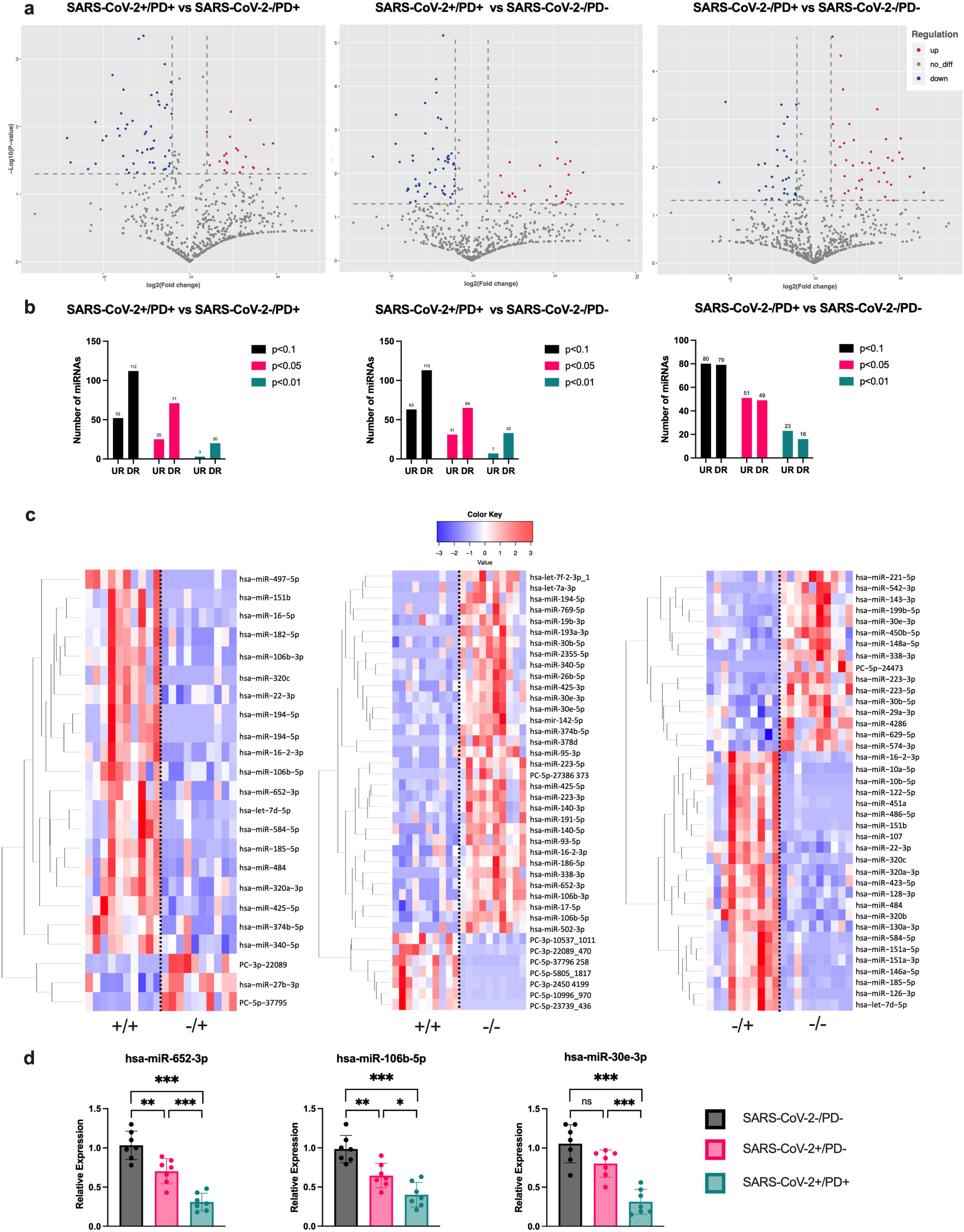
Dysregulation of miRNA Expression in SARS-CoV-2+/PD+ (LC) vs SARS-CoV-2-/PD+ Subjects. (**a)** Volcano plot representing differential salivary miRNA expression in subjects with or without PD and/or a history of COVID-19 (n= 10/group). SARS-CoV-2-/PD+ (Group 1); SARS-CoV-2+/PD+ (LC) (Group 2); SARS-CoV-2-/PD-(Group 3). (**b)** Quantification of differential salivary miRNA expression across the different groups. (**c)** Heatmap of differential miRNA expression in Group 2 vs Group 3 (p<0.01). For paired comparisons, the significance of differential expressions of detected miRNAs was assessed using Student’s t-test on normalized expression values. Hybridization signals were normalized prior to analysis (+/+: SARS-CoV-2+/PD+ (LC); −/+: SARS-CoV-2-/PD+; −/−: SARS-CoV-2-/PD-). Validation of selected dysregulated miRNAs in a post-vaccination cohort by RT-qPCR. Total RNA was isolated from saliva samples (n=7/group) obtained from study participants and miRNA expression was quantified by RT-qPCR. Bar graphs showing relative expression of miR-30e-5p (**d**), miR-106b-5p (**e**), and miR-652-3p (**f**) in an independent post-vaccination cohort showed expression patterns consistent with those observed in the pre-vaccination sequencing dataset. Statistical significance was determined using one-way ANOVA with multiple comparisons, *\*\*\*P < 0.001, **P < 0.01, *P < 0.05* were considered significant. ns = not significant.

To validate the sequencing-derived miRNA signature, we assessed the expression of the selected dysregulated miRNAs miR-30e-3p, miR-106-5p and miR-652-3p in an independent post-vaccination cohort using RT-qPCR. Compared to pre-vaccination cohort, a similar expression pattern was observed across the validated candidates. Expression of miR-30e-3p (fold change: 0.32 ± 0.11), miR-106-5p (fold change: 0.41 ± 0.15), and miR-652-3p (fold change: 0.3 ± 0.17) were markedly reduced in SARS-CoV-2+/PD+ compared to SARS-CoV-2-/PD+ (fold change: 0.71 ± 0.16; 0.64 ± 0.15; 0.8 ± 0.15) or SARS-CoV-2-/PD- (**Fig.1 D-F**). This indicates that the post-vaccination cohort also exhibit similar miRNA expression profiles observed in pre-vaccination miRNA sequencing.

### Dysregulated miRNAs In SARS-CoV-2+/PD+ (LC) Patients Regulate Host Inflammatory Response

One mechanism by which host miRNAs influence viral infection is by binding directly to the microRNA response element (MRE) sites of host mRNAs as demonstrated in prior studies.^36–39^ This interaction fine-tunes the expression of immune-related genes at the post-transcriptional level, thereby suppressing the antiviral immune response. The Kyoto Encyclopedia of Genes and Genomes (KEGG) pathway analysis of genes regulated by differentially expressed miRNAs revealed significant enrichment in pathways related to cell signaling, immune responses, and cancer progression (**Fig.2A**). Pathways were ranked based on their rich factor, gene count, and p-value, providing insights into the biological processes influenced by the differentially expressed miRNAs. Among the most enriched pathways was the Renin-angiotensin system (Ras), which displayed a high gene count and a rich factor nearing 0.98 (**Fig.2A**). Given the well-established role of the Ras pathway in facilitating SARS-CoV-2 entry and mediating inflammation, this pathway was prioritized for subsequent analysis.^40,41^ Moreover, due to the canonical inverse relationship between miRNA and mRNA expression, we focused on the downregulated miRNAs in SARS-CoV-2+/PD+ (LC) subjects, hsa-miR-106b-3p, hsa-miR-106b-5p, hsa-miR-652-3p, and hsa-miR-30e-3p.

**Fig. 2.**
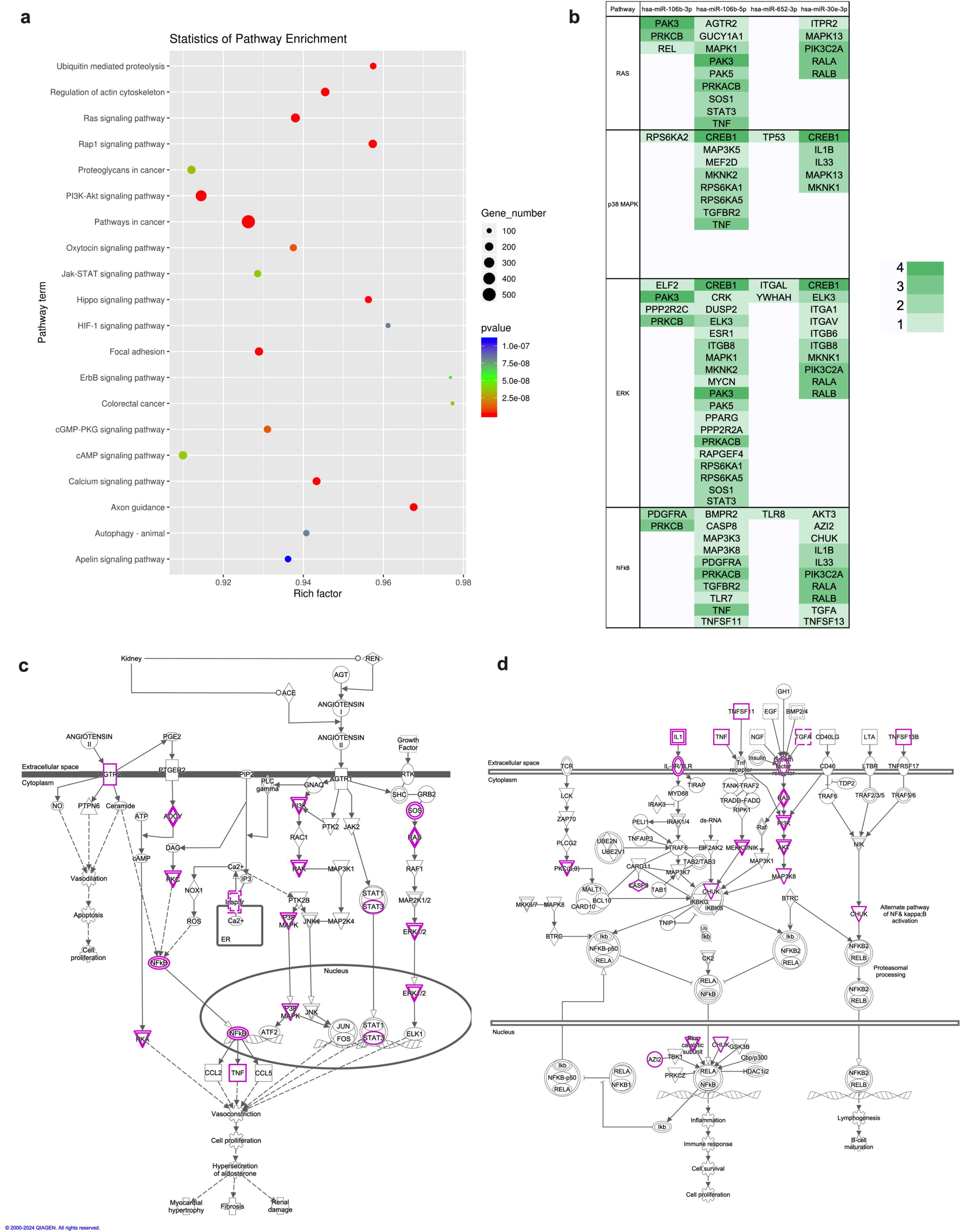
Enrichment Analysis and Target Prediction of Differentially Expressed miRNAs. (**a)** Scatterplot of enriched KEGG pathways showing the rich factor and p-value for various signaling pathways. The rich factor (calculated as the number of target genes in a pathway divided by the total number of genes in that pathway) indicates the level of enrichment, with larger values representing higher enrichment. The size of each circle corresponds to the number of genes involved in each pathway, while the color reflects the p-value. (**b)** Ingenuity Pathway Analysis (IPA) showing the target genes of hsa-miR-106b-3p, hsa-miR-106b-5p, hsa-miR-652-3p, and hsa-miR-30e-3p categorized by RAS and its downstream signaling pathways (p38 MAPK, ERK, and NFkB). Green shading indicates genes with significant involvement in these pathways, with darker shades representing higher incidence (1-4). Target genes of hsa-miR-106b-3p, hsa-miR-106b-5p, hsa-miR-652-3p, and hsa-miR-30e-3p in the (**c)** RAS, (**d)** IL-1β, and TNF pathways are highlighted in purple.

To determine which of the specific genes within the Ras pathway are targets of our candidate miRNAs, we conducted an Ingenuity Pathway Analysis (IPA). This analysis identified 17 putative immune gene targets within the Ras pathway regulated by miR-106b-3p, miR-106b-5p, and miR-30e-3p (**Fig.2B-C**). However, our analysis failed to identify any genes directly under Ras for miR-652-3p (**Fig.2B**). Because Ras is a central signaling hub with multiple downstream effectors, we further used IPA to isolate its inflammatory signaling components. Among the Ras-influenced inflammatory cascades targeted by the candidate miRNAs, we focused on the mitogen-activated protein kinases (MAPK) p38 and ERK, as well as NFκB, three pathways implicated in COVID-19-driven hyperinflammation (**Fig.2B-C**).^42–44^ IPA analysis reveals that p38, ERK, and NFκB pathways share a multitude of inflammation-modulatory genes, of which we highlight IL-1β and TNFs (TNF□, TNFSF11, and TNFSF13B) (**Fig.2B-D**). IL-1β and TNFs are not only well-known regulators of inflammation but are central to the pathobiology of PD and COVID-19.^45^ The overlapping regulation of these signaling cascades by miR-106b-3p, miR-106b-5p, and miR-30e-3p suggests a mechanism by which miRNA downregulation may contribute to the heightened inflammatory state observed in both diseases.

### Downregulation of miR-30b-3p, miR-106b-3p, and miR-106b-5p potentiates innate immune activation in oral epithelium

To define the role of miR-30e-3p, 106b-3p, and miR-106b-5p in regulating inflammatory responses, gingival keratinocytes were transfected with miRNA mimics, inhibitors, or their respective controls and subsequently challenged with the TLR4 agonist *E. coli* LPS (50 ng/mL). NFκB activation was evaluated by flow cytometric measurement of phosphorylated NFκB (p-NFκB), and cytokine release was quantified in culture supernatants collected 12 h after stimulation. Histograms show that in cells transfected with miR-30e-3p, 106b-3p, and miR-106b-5p mimics, TLR4 stimulation induced substantially lower levels of p-NFκB than in control mimic-transfected cells (**Fig.3A**). In contrast, transfection of miR-30e-3p, miR-106b-3p, or miR-106b-5p inhibitors enhanced the inflammatory response to TLR4 stimulation. Cells transfected with miRNA inhibitors exhibited increased p-NFκB compared with inhibitor control-transfected cells (**Fig.3A**). Quantification using control normalized geometric mean fluorescence intensity (geo-MFI) confirmed a significant reduction in p-NFκB in mimic-transfected cells expressing miR-30e-3p (64.29 ± 11.12), 106b-3p (79.55 ± 2.72), and miR-106b-5p (58.91 ± 8.48), while miRNA inhibitors exhibited antagonistic function as observed by markedly higher expression of p-NFκB [miR-30e-3p (127.3 ± 5.1), 106b-3p (120.3 ± 6.6), and miR-106b-5p (118.7 ± 7.7)] (**Fig.3B**). These findings suggest that downregulated miRNAs act as negative regulators of inflammatory signaling by suppressing NFκB activation.

**Fig. 3.**
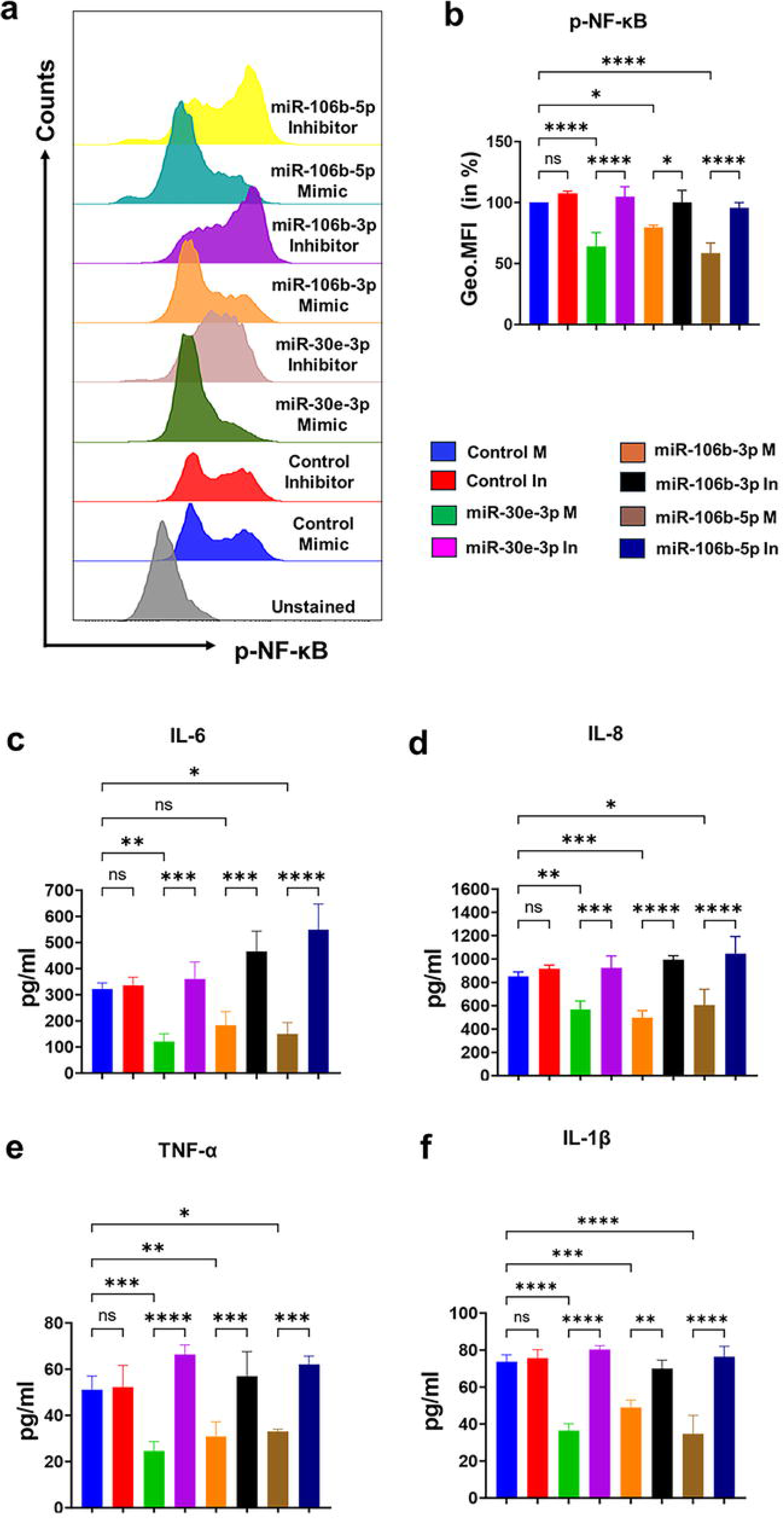
miR-30e-3p, miR-106b-3p, and miR-106b-5p suppress TLR4-driven inflammatory activation in oral keratinocytes. Primary gingival keratinocytes were transfected with miRNA mimics, inhibitors, or corresponding controls (25 nM) using Lipofectamine 2000. Cells were stimulated with TLR4 agonist *E. coli* LPS (50 ng/mL). NFκB activation was assessed by flow cytometric detection of phosphorylated NFκB (p-NFκB), and cytokine secretion was measured in culture supernatants collected 12 h after stimulation. (**a)** Representative flow cytometry histogram showing p-NFκB expression in LPS-stimulated gingival keratinocytes. (**b)** Bar graphs showing normalized geometric mean fluorescence intensity (geo. MFI) values expressed as percentages relative to control mimic. Data is presented as mean ± SEM from three replicates. Multiplex ELISA was performed to quantify cytokines in the supernatant. Bar graphs show levels of pro-inflammatory cytokines (**c**) IL-6, (**d**) IL-8, (**e**) TNF-α, and (**f**) IL-1β (pg/ml) quantified by multiplex bead array. Data are presented as mean ± SEM from four independent donors. Statistical significance was determined by unpaired two-tailed Student’s t-test. p < 0.05 was considered significant.

We next investigated whether the observed attenuation of NFκB activation correlated with cytokine production. Oral keratinocytes were transected with miRNA mimics or inhibitors and challenged with *E. coli* LPS. Cells expressing miR-30e-3p, miR-106b-3p, or miR-106b-5p mimic exhibit markedly lower supernatant levels of IL-1β (36.33 ± 3.78 pg/ml; 49.06 ± 4.77 pg/ml; 34.92± 10.53 pg/ml), TNF-α (24.18 ± 4.04 pg/ml; 30.86 ± 7.73 pg/ml; 33.34 ± 1.64 pg/ml), IL-6 (120.90 ± 30.66 pg/ml; 183.78 ± 52.84 pg/ml and 150.19 ± 43.58 pg/ml), and IL-8 (570.93 ± 69.87 pg/ml; 496.99 ± 61.10 pg/ml; 606.66 ± 134.28 pg/ml) compared with control mimic transfected cells (**Fig.3C-F**). In contrast, inhibition of these miRNAs increased secretion of all four cytokines relative to corresponding mimics and inhibitor controls (**Fig.3C-F**). These findings demonstrate that miR-30e-3p, miR-106b-3p, and miR-106b-5p suppress TLR4-induced pro-inflammatory cytokine production and function as endogenous anti-inflammatory regulators in oral keratinocytes.

### Downregulated miRNAs in SARS-CoV-2+/PD+ (LC) Patients Target SARS-CoV-2 Spike and Nucleocapsid Transcripts

Drawing from our miRNA-seq data and previous reports demonstrating that global miRNA repression is a hallmark of many diseases ^46^, we focused our subsequent analysis on known host miRNAs that were significantly downregulated in SARS-CoV-2+/PD+ (LC) patients compared to healthy controls (p < 0.01) (**Table 1**). To investigate how these candidate miRNAs might influence SARS-CoV-2 pathogenesis, we performed bioinformatic analysis to identify those with putative binding sites on viral RNA transcripts encoding the structural proteins spike and nucleocapsid. Our previous *in silico* analysis revealed that: (1) a substantial proportion of all known mature human miRNAs (ranging from 598 to 1,229 out of 2,636 miRNAs) have predicted binding sites across various SARS-CoV-2 genomes; and (2) while minimal differences in miRNA binding were observed among earlier - Wuhan-Hu-1 (1,229), Beta (B.1.351: 1,227 miRNAs), and Delta (B.1.617.2: 1,232 miRNAs) - a marked reduction in predicted human miRNA interactions was noted in Omicron (BA.1: 598 miRNAs) ^31^. Therefore, of all the SARS-CoV-2 variants, we focused on Beta (B.1.351) and Omicron (BA.1) for further analysis. Beta was selected due to its representative miRNA-binding profile among early variants and its well-defined spike mutations that enable mechanistic interrogation of miRNA-mediated effects on viral entry.

Using stringent binding parameters (energy threshold=-25 kcal/mol), we identified four spike-targeting miRNAs, miR-652-3p, miR-425-5p, miR-106b-3p, and miR-106b-5p, all of which were predicted to bind both the Beta and Omicron spike sequences (**Fig. 4A-B**). *In silico* analysis further revealed that these miRNAs target key spike regions, including the N-terminal domain (NTD), receptor-binding domain (RBD), and the SD1-S2 region (**Fig.S1**). In contrast, these computational parameters failed to predict any miRNA:nucleocapsid interactions. After relaxing the energy threshold to −10 kcal/mol, we identified nine nucleocapsid-targeting miRNAs. Among these nucleocapsid-targeting miRNAs, eight were predicted to target both variants: let-7a-3p, miR-2355-5p, miR-30e-3p, miR-378d, miR-223-5p, miR-223-3p, miR-140-5p, and miR-652-3p (**Fig.4C)**. In contrast, miR-26b-5p uniquely targeted the Omicron nucleocapsid transcript. Pairwise comparison analysis revealed that miR-652-3p was the only miRNA predicted to target both the spike and nucleocapsid transcripts across both variants.

**Fig. 4.**
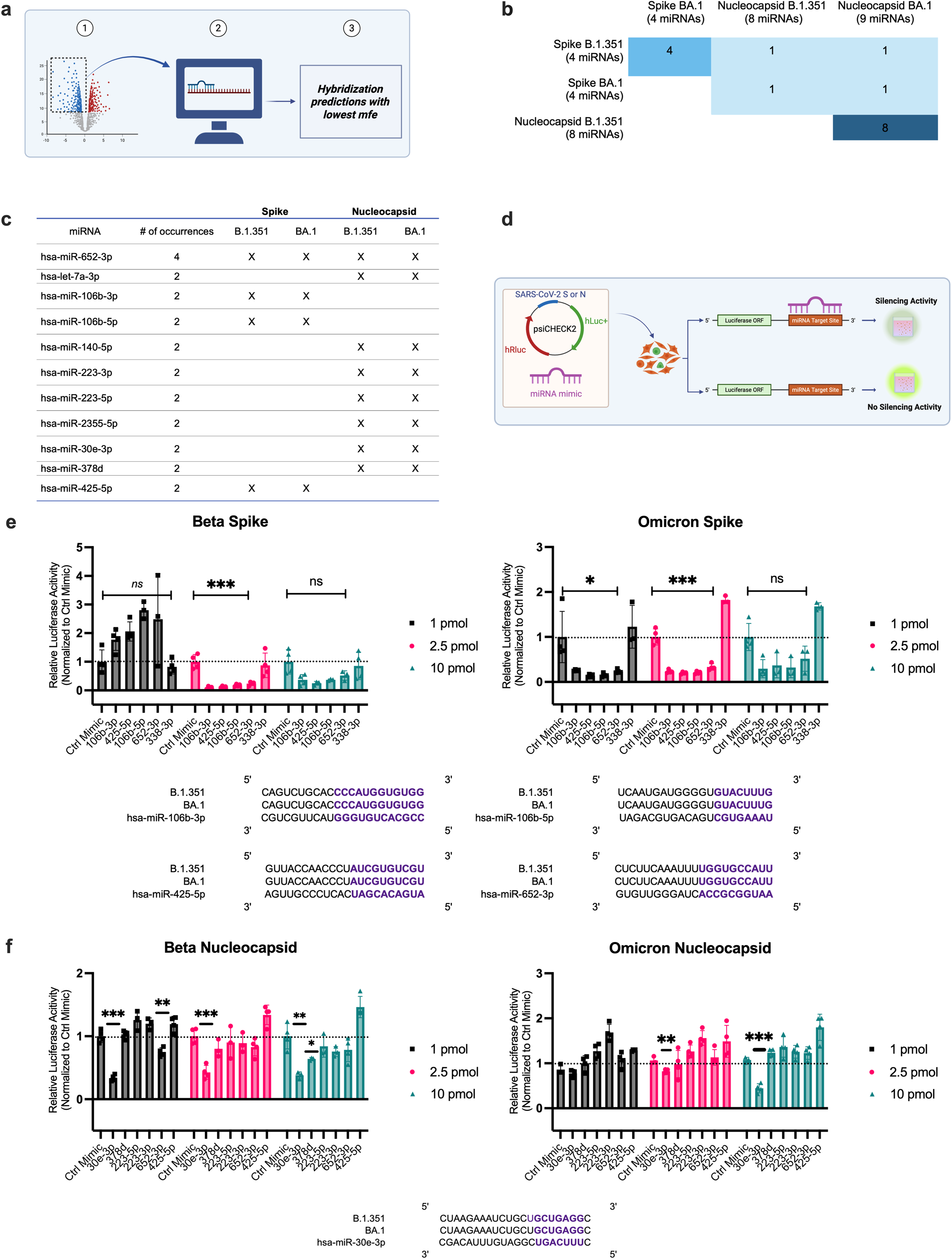
Identification and validation of host microRNAs predicted to target SARS-CoV-2 Spike and Nucleocapsid sequences. **(a)** Schematic overview of the computational pipeline used to identify host miRNAs predicted to bind viral sequences based on hybridization energy (RNAhybrid 2.0), selecting candidates with the lowest minimum free energy (mfe). (**b)** Pairwise comparison matrix and (**c)** Table showing the overlap of predicted miRNAs targeting B.1.351 and BA.1 Spike and Nucleocapsid regions. (**d)** Schematic of dual-luciferase reporter assay used to verify miRNA and viral transcript interactions. SARS-CoV-2 Spike or Nucleocapsid sequences (B.1.351 or BA.1 variants) were subcloned into the psiCHECK-2 dual-luciferase reporter vector and co-transfected with miRNA mimics into HEK293T cells. The target gene is cloned downstream of a luciferase reporter gene. This includes the predicted binding site of the miRNA. If the miRNA being tested can directly bind to the complementary sequence in the reporter construct, it results in reduced expression of luciferase and a decrease in luminescence. (**e)** Relative luciferase activity (normalized to control mimic) in co-transfected HEK293T cells at increasing concentrations of miRNA mimics. A value of relative luciferase activity <1 indicates effective miRNA binding to either B.1.351 or BA.1 Spike or (**f)** Nucleocapsid, with binding site alignment sequences shown below. Data represent mean±SD from biological replicates (n = 4). Statistical significance was determined using one-way ANOVA with multiple comparisons; *\*\*\*\*P < 0.0001*, *\*\*\*P < 0.001, **P < 0.01, *P < 0.0*5 were considered significant. ns = not significant.

Next, to validate these predicted miRNA:spike and miRNA:nucleocapsid interactions we performed dual luciferase reporter assays. The direct binding of a miRNA to its target sequence reduces luciferase signal (**Fig. 4D**). Luciferase assay results showed significant suppression of relative luciferase activity in cells overexpressing miR-652-3p, miR-425-5p, miR-106b-3p, and miR-106b-5p (2.5 pmol of miRNA mimic) indicating direct regulation of Beta and Omicron spike transcript (p<0.001) (**Fig. 4E**). Similarly, we quantified consistent and reduced relative luciferase activity upon the binding of miR-30e-3p (2.5 pmol and 10 pmol of miRNA mimic) to Beta and Omicron nucleocapsid (**Fig. 4F**). In addition to the mirVana^TM^ control mimic (Invitrogen, Waltham, MA), miR-338-3p and hsa-miR-425-5p served as additional negative controls for spike and nucleocapsid, respectively, and were selected based on our bioinformatic analysis. We highlight that neither hsa-miR-338-3p nor hsa-miR-425-5p were experimentally validated to significantly target spike or nucleocapsid, respectively, showcasing the robust efficacy of our in-silico analysis (**Fig. 4E-F**). Altogether, these findings confirm the direct binding of select miRNAs to SARS-CoV-2-spike and nucleocapsid sequences.

### miR-30e-3p, miR-106b-3p, and miR-652-3p reduces SARS-CoV-2 spike protein and nucleocapsid transcript expression

After confirming direct targeting of SARS-CoV-2 spike transcripts by hsa-miR-106b-3p, hsa-miR-425-5p, hsa-miR-106b-5p, and hsa-miR-652-3p, and nucleocapsid transcripts by hsa-miR-30e-3p, we further investigated the role of host-encoded miRNAs in modulating SARS-CoV-2 pathogenesis. Host miRNAs have previously demonstrated antiviral capability by binding directly to viral transcripts and subsequently degrading or suppressing (1) viral RNA or (2) the translation of proviral proteins.^47–49^ Considering this miRNA-mediated antiviral mechanism and the established roles of spike and nucleocapsid proteins in facilitating SARS-CoV-2 pathogenesis, we hypothesized that miRNA:spike and miRNA:nucleocapsid interactions may attenuate SARS-CoV-2 replication.

To test our hypothesis, we transfected SARS-CoV-2-permissive ACE2 expressing A549 epithelial cells with miRNA mimics or overexpression plasmids corresponding to miR-30e-3p, miR-106b-3p, and miR-652-3p–three miRNAs that were among the most significantly downregulated (i.e., largest negative log2 fold change) in +PD/+SARS-CoV-2+ versus SARS-CoV-2-/PD- patients, and that were experimentally confirmed to target SARS-CoV-2 spike (miR-106b-3p and miR-652-3p) or nucleocapsid (miR-30e-3p) transcripts (**Table 1**). Following transfection, cells were challenged with either the SARS-CoV-2 Beta (B.1.351) or Omicron (B.1.1.529) variant, and spike and nucleocapsid transcript expression levels were quantified at 24 or 48 hours post-infection (hpi) via RT-qPCR and flow cytometry, respectively. Flow cytometry analysis revealed significant reductions in Spike-positive cells following transfection with all three miRNA mimics across both variants and time points (**Fig. 5A-B, E-F**). The effect was most pronounced at 24 hpi, suggesting that these miRNA mimics act early in the viral life cycle (**Fig. 5A-B, E-F**). Among the mimics at 24hpi, miR-106b-3p consistently exhibited the strongest suppression, reducing Spike-positive cells by 84% and 78% in B.1.351- and B.1.1.529-infected cells, respectively, compared to control (**Fig. 5A-B, E-F**). Similar trends were observed with pCMV-based miRNA expression (**Fig. 5C-D, G-H**), although the suppressive effect was generally less potent than that of the corresponding mimics. Additionally, the suppressive effect of pCMV-652 diminished by 48 hpi in B.1.1.529-infected cells, suggesting that delivery strategy influences the durability of spike suppression (**Fig. 5G-H**).

**Fig. 5.**
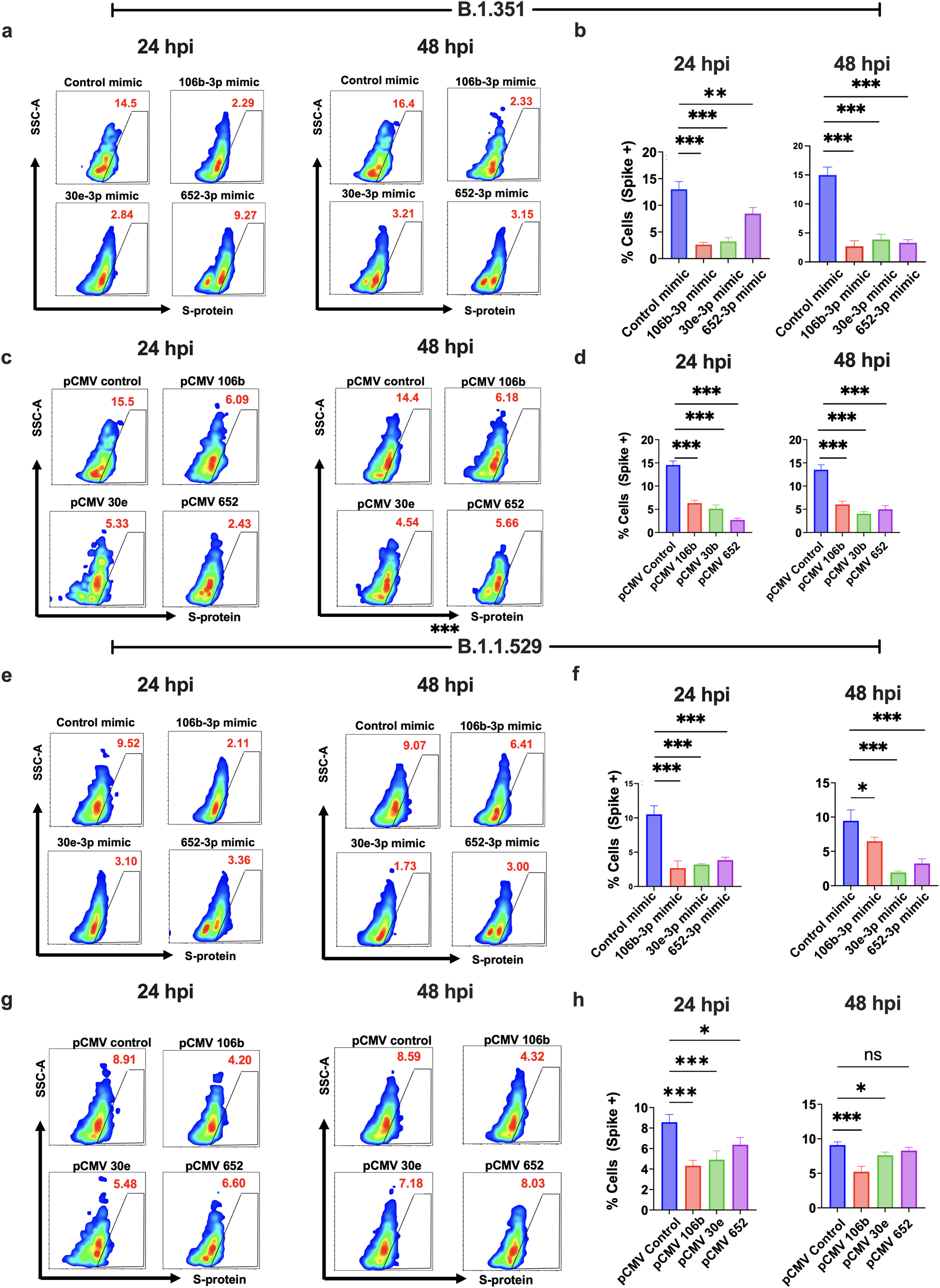
MicroRNA overexpression reduces SARS-CoV-2 spike protein expression. Cells were transfected with miRNA mimics or overexpression plasmids and infected with B.1.351 or B.1.1.529 strains. (**a)** Representative flow cytometry scatter plots showing spike protein-positive cells at 24 and 48 hpi with either miRNA mimics (**a**, **e**) or overexpression plasmids (**c**, **g**), followed by exposure to B.1.351 (**a-d**) or B.1.1.529 (**e-h**). **b, d, f, h** Corresponding quantification of spike-positive cells from flow cytometry data at 24 and 48 hpi. Data represents SD from biological replicates (n=4). Statistical significance was determined using one-way ANOVA with multiple comparisons, *\*\*\*\*P < 0.0001, ***P < 0.001, **P < 0.01, *P < 0.05* were considered significant. ns = not significant.

The dichotomy between miRNA delivery via mimic versus plasmid became more pronounced when assessing SARS-CoV-2 replication by quantifying nucleocapsid (N) gene expression. While flow cytometry showed consistent suppression of Spike protein across both delivery methods, mimic-based delivery resulted in more potent and sustained inhibition of viral RNA levels. Specifically, RT-qPCR analysis revealed an approximately ten-fold reduction in nucleocapsid expression in miR-652-3p-transfected cells at 48 hpi with B.1.351 (p < 0.01), which was also observed in B.1.1.529-infected cells at both 24 hpi (p < 0.01) and 48 hpi (p < 0.0001). In contrast, miR-30e-3p transfection decreased nucleocapsid expression only at 24 hpi (∼2.4 fold) with B.1.351 (p < 0.01), but not at 48 hpi or in B.1.1.529-infected cells (**Fig. 6A-B**). Nucleocapsid expression significantly increased in miR-106b-3p-transfected cells 48 hpi with B.1.1.529 (∼1.36 fold) (**Fig. 6C**). In contrast, the corresponding pCMV constructs yielded only transient or minimal effects (**Fig. 6B, D**). At 24 hpi with B.1.1.529, transfection with pCMV-30e and pCMV-106b led to modest reductions in N gene copies (p<0.05 to p<0.01), but these effects were not sustained at 48 hpi (**Fig. 6D**). Moreover, nucleocapsid expression also increased in hsa-652-transfected cells 24 hpi with B.1.351 (**Fig. 6B**). Overall, miRNA mimics exhibit more robust and sustained reductions in both Spike and nucleocapsid RNA expression levels in both variants at both 24 and 48 hpi.

**Fig. 6.**
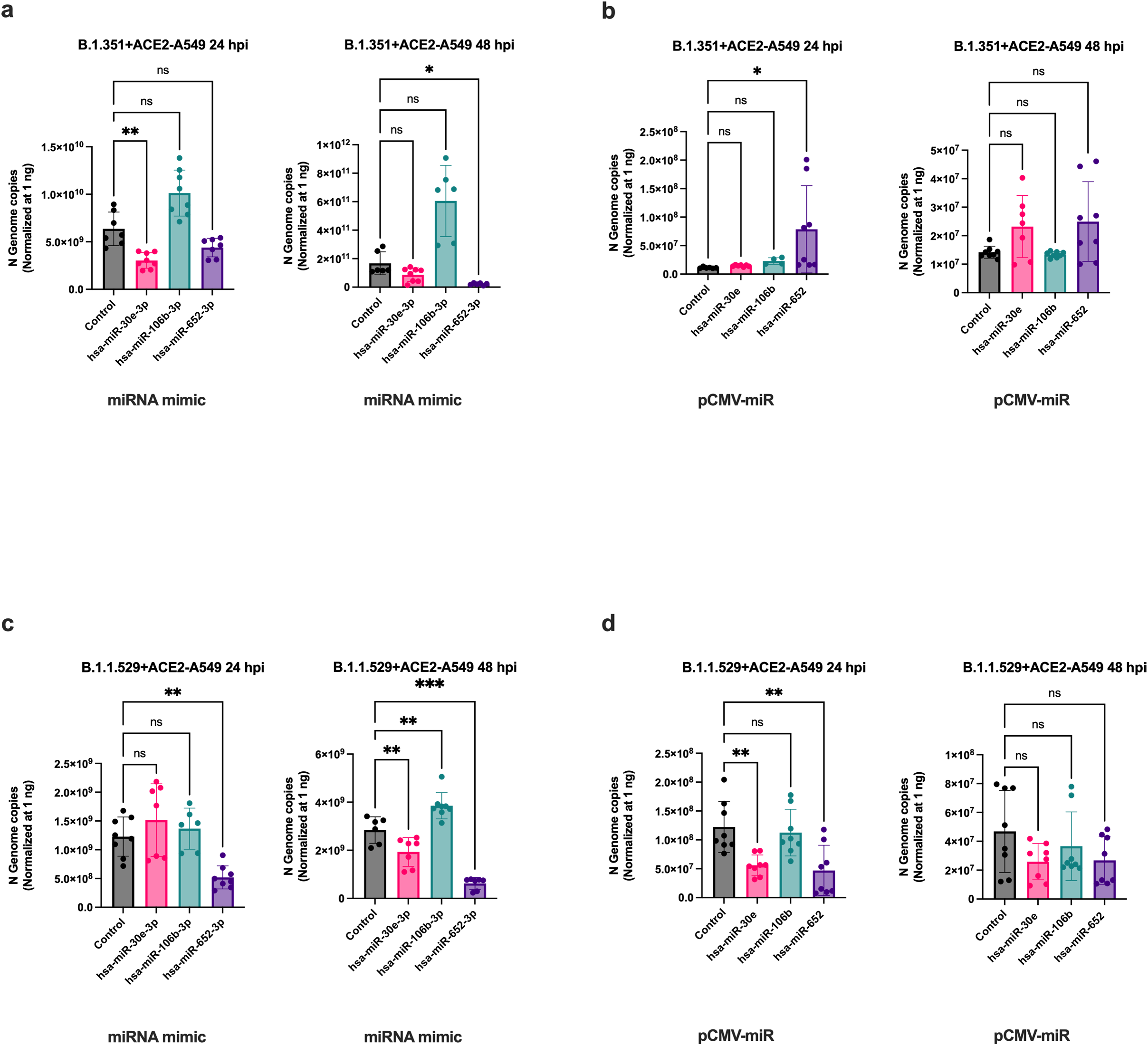
miRNA overexpression reduces SARS-CoV-2 replication. ACE2-A549 cells were transfected with miRNA mimics or overexpression plasmids and infected with SARS-CoV-2. Viral [Nucleocapsid (N)] genome copies following transfection of ACE2-A549 cells with (**a)** miRNA mimics and (**b)** overexpression plasmids at 24 and 48 hpi (B.1.351). c-d N genome copies following transfection of ACE2-A549 cells with (**c)** miRNA mimics and (**d)** overexpression plasmids at 24 and 48 hpi (B.1.1.529). All data represent the mean ± SD from 4-5 biological replicates. Statistical significance was determined using one-way ANOVA with Kruskal-Wallis test. * ns = not significant.

## DISCUSSION

Our study explores the interplay between PD and SARS-CoV-2 infection, emphasizing their converging mechanisms of inflammation and viral persistence within the oral cavity. Previous clinical examination of our cohort revealed that regardless of vaccination status COVID-19 history correlates with worsened periodontal health, as evidenced by increased probing depths, plaque index, gingival index, and bleeding index—all markers of heightened periodontal inflammation.^14–17^ In the same patients, RT-qPCR analysis confirmed elevated salivary expression of proinflammatory mediators, IL-6, IL-8, and matrix metalloproteinase 8 (MMP-8), as well as the SARS-CoV-2 entry receptors ACE2 and TMPRSS2 (Schwartz et al. Unpublished findings).^17^

The regulatory mechanisms driving this immune axis in Long COVID remain poorly understood, but growing evidence suggests that viral persistence may be an important contributing factor. Persistent viral reservoirs or residual viral antigens could continuously stimulate the immune system, thereby perturbing immune regulation and promoting chronic inflammation. The underlying mechanisms linking viral persistence to this immune axis remain largely understudied. We demonstrate unique perturbations in the salivary miRNA profiles of SARS-CoV-2+/PD+ subjects, suggesting a distinct molecular signature driven by the intersection of SARS-CoV-2 infection and periodontal disease, in comparison with SARS-CoV-2+ subjects without PD, thereby highlighting PD-associated modulation of viral infection–specific miRNome changes. Consistent with our findings, Saulle et al. identified significant downregulation of salivary and circulating let-7a-5p, let-7b-5p, and let-7c-5p in patients with acute COVID-19 infection, whereas production of pro-inflammatory cytokines and chemokines, such as IL-1β, IL-6, IL-8, and TNF-□, increased in the saliva and plasma of mild and severe acute COVID-19 patients.^50^ These findings underscore the potential for sustained immune activation and viral activity in the oral cavity, particularly among individuals with concurrent PD and acute or post-acute COVID-19 implying role of viral persistence in dysregulating oral mucosal immunity.

The global downregulation of salivary miRNAs observed in SARS-CoV-2+/PD+ (LC) patients further support the role of miRNAs in maintaining immune homeostasis. Widespread miRNA repression has been previously associated with systemic immune-mediated conditions and is increasingly viewed as a hallmark of immune dysregulation.^45,51^ In our cohort, downregulated miR-106b-3p, miR-106b-5p, and miR-30e-3p target components of Ras-associated inflammatory signaling, including IL-1β and TNF- family members, both implicated in the pathogenesis of PD and COVID-19. In PD, proinflammatory cytokines like TNFα recruit neutrophils and macrophages to the periodontium, where they release MMPs and reactive oxygen species (ROS), contributing to tissue degradation and bone loss. Similarly, in COVID-19, dysregulated immune responses can escalate into a cytokine storm, a severe hyperinflammatory state marked by widespread cytokine release and multi-organ injury. In fact, miR-30e-3p is reportedly downregulated in acute COVID-19+ patients with cardiovascular disease, supporting the miRNA’s potential contribution to the inflammatory response.^52^ These findings suggest that miRNA downregulation may exacerbate inflammation in both diseases through shared molecular pathways.^53^

Beyond their role in immune regulation, host miRNAs may exert antiviral effects by targeting key steps in the SARS-CoV-2 replication cycle–a concept supported by our and other groups’ data.^54–57^ For example, transfection of serum-isolated miR-223-3p, which we also identified as downregulated in SARS-CoV-2+/PD+ (LC) subjects compared to healthy controls (**Table 1**), into HEK293T cells decreased spike mRNA levels.^53^ Similarly, our observed suppression of spike protein expression following miRNA mimic transfection suggests a potential interference with viral entry (**Fig.4**), considering the spike protein’s essential role in ACE2 binding and membrane fusion. In fact, *in silico* analysis further predicted that hsa-miR-30e-3p specifically targets spike at the RBD (**Fig.S1**). It is well-documented that the SARS-CoV-2 spike engages ACE2 on host cells via the viral protein’s RBD. This interaction is followed by proteolytic cleavage of spike at the S1/S2 and/or S2’ sites by host proteases, enabling viral and host membrane fusion.^31,58^ We highlight that the strongest reductions occurred at 24 hpi, supporting the hypothesis that these miRNAs act robustly during initial infection, potentially limiting viral dissemination before robust replication is established. However, further analysis using viral entry assays is required to substantiate the role of these candidate miRNAs in viral entry. Nonetheless, by downregulating spike expression, these host miRNAs may serve as intrinsic antiviral factors that restrict viral spread during the early stages of infection. Given the high mutational burden of the spike gene across SARS-CoV-2 lineages, the ability of our candidate miRNAs to retain antiviral activity across variants supports their potential as mutation-resilient therapeutic tools.^59–61^

Targeting and subsequent suppression of nucleocapsid transcripts by miR-652-3p and miR-30e-3p reveals an additional layer of antiviral activity during the replication phase. Nucleocapsid plays a critical role in viral RNA packaging, replication complex, assembly, and genome stability, making it indispensable for productive infection.^62^ Suppression of nucleocapsid transcript may thereby directly interfere with virion production, although further experimentation is needed for confirmation.

Our data also underscores the dual potential of host miRNAs to either restrict or assist viral replication. For instance, nucleocapsid RNA levels increased in miR-106b-3p transfected cells at 48 hpi with B.1.1.529 and in miR-652 transfected cells at 24 hpi with B.1.351. These increases may reflect off-target effects, a well-documented limitation of miRNA-based therapeutics. Although miRNA mimics and plasmid-based expression systems are designed to act in a target transcript-specific manner, neither approach guarantees strict specificity in practice.^63^ Off-target effects can arise because miRNAs require only partial complementarity between their seed region and target mRNAs to exert regulatory effects; they are inherently capable of binding to multiple unintended transcripts.^64^ This promiscuity is exacerbated by the high concentrations typically used in transfection protocols, such as the 50 nM dose employed in our mimic experiments.^65^ The differential effects we observed- strong and consistent suppression of spike protein versus more variable effects on nucleocapsid transcript levels- further support this model. Partial complementarity tends to inhibit translation, whereas perfect or near-perfect base pairing is more likely to trigger mRNA degradation.^66^ Thus, differences in miRNA-target pairing fidelity across spike and nucleocapsid transcripts may account for the variable efficacy and occasional paradoxical upregulation observed in our system.

The mode of miRNA delivery into cells may also account for the discrepancies in our data, underscoring the importance of dosage, timing, and subcellular localization in achieving effective silencing-factors that are often overlooked in miRNA-based antiviral strategies. miRNA mimics and plasmids are used to introduce or elevate levels of specific miRNAs in cells, albeit through distinct mechanisms: mimics supply mature, double-stranded RNA delivery to the cytoplasm for immediate incorporation into RNA-induced silencing complex (RISC), whereas plasmids rely on transcription and endogenous processing through the canonical miRNA biogenesis pathway.^67,68^ Thus, the transient suppression observed with plasmid-based expression, particularly in Omicron-infected cells, may reflect delayed processing and lower levels of viral transcript-targeting mature miRNAs from plasmid-based transcription compared to direct cytoplasmic delivery of mimics. Altogether, our findings suggest that while both mimic- and plasmid-based delivery can transiently suppress Spike protein expression, mimic-based delivery more effectively and durably inhibits SARS-CoV-2 replication reflected by sustained reductions in both surface spike and intracellular nucleocapsid RNA.

Although our experimental work focused on earlier SARS-CoV-2 variants, including Beta and Omicron subvariants BA.1 and B.1.1.529, a separate *in silico* analysis confirms that miR-106b-3p, miR-30e-3p, and miR-652-3p retain binding potential in the more recent XFG genome (**Fig.S2**), supporting their continued therapeutic relevance. While these results highlight the potential of miRNAs as modulators of inflammation and viral activity, the upstream mechanisms driving their dysregulation remain unclear and merit further investigation.

While other groups have previously achieved systemic and salivary miRNA profiling in COVID-19 patients, particularly those with acute infection, we highlight miRNA changes and provide potential biomarkers in long COVID patients with concomitant periodontitis. Crucially, there is a lack of salivary biomarkers for this patient population, and this deficit underscores the urgent need for accessible and non-invasive diagnostic tools given the prevalence of long COVID and its effect on systemic health, oral-systemic interactions, and overall quality of life. To this end, we emphasize three host miRNAs–miR-652-3p, miR-106b-3p, and miR-30e-3p–as candidates, given their promising antiviral and inflammation-modulating properties. Interestingly, miR-30e-3p had previously demonstrated the capability of inhibiting influenza B virus (ssRNA) replication by targeting the virus’ nucleoprotein (NP) and neuraminidase (NA) genes.^69^ Furthermore, KEGG pathway enrichment of Ras, NFκB, and MAPK signaling reinforces the systemic implications of miRNA loss and points to candidate pathways for intervention. Studies from our lab have demonstrated that miR-30 family members exert anti-inflammatory activity by negatively regulating immune activation. In myeloid inflammatory cells (macrophages or dendritic cells) challenged with *E. coli* or IgG, miR-30b suppress inflammatory signaling through downregulation of pathways associated with macrophage activation, thereby limiting innate immune responses, while also constraining adaptive immune activity.^70–72^ Together, these findings highlight the miR-30 family as an important regulator of immune homeostasis. Taken together, these findings provide a novel framework for understanding miRNA-mediated crosstalk between PD and long COVID and identify actionable targets for future therapeutic strategies aimed at mitigating inflammation and improving patient outcomes.

In conclusion, these data uncover a previously unrecognized miRNA-dependent mechanism by which host immune dysregulation may drive viral persistence in the oral compartment during Long COVID. In SARS-CoV-2-positive individuals with PD, broad repression of salivary miRNAs contributes to both immune hyperactivation and failure of viral clearance. By demonstrating that restoration of specific miRNAs restrains NFκB-driven cytokine responses while directly suppressing SARS-CoV-2 transcripts, our study demonstrates a key role of host miRNA as SARS-CoV-2 restriction factor and oral mucosal immune activity. This work therefore defines a mechanistic intersection between periodontal inflammation and persistent SARS-CoV-2–associated pathology, with important implications for biomarker development and miRNA-based therapeutic intervention.

## AUTHOR CONTRIBUTIONS

K.J.C. and A.R.N conceived and designed the study. K.J.C., R.A.N., S.E., S. E., and J.C. conducted most experiments. K.J.C. ran and analyzed all computational analyses. K.J.C., R.A.N., and A.R.N. conducted most data analysis. A.R.N, J.L.S., and J.R. co-supervised the study. K.J.C. and A.R.N wrote the manuscript, which the other authors edited and approved.

## CONFLICT of INTERESTS

The authors declare no competing interests.

## FUNDING

This study was supported by Contract grant sponsor: NIDCR/NIH; contract grant numbers: R01DE027980 (ARN), R56DE033249 (ARN and JS), and NEI/NIH contract grant number R01EY033622 (ARN).

## ABBREVIATIONS

ACE2: Angiotensin-converting enzyme 2
ANOVA: Analysis of variance
BA.1: Omicron subvariant BA.1
BSA: Bovine serum albumin
Ct: Cycle threshold
COVID-19: Coronavirus disease 2019
ERK: Extracellular signal-regulated kinase
GISAID: Global Initiative on Sharing All Influenza Data
HIPAA: Health Insurance Portability and Accountability Act
HOK: Human oral keratinocytes
IL: Interleukin
IPA: Ingenuity Pathway Analysis
IRB: Institutional Review Board
KEGG: Kyoto Encyclopedia of Genes and Genomes
LC: Long COVID
LPS: Lipopolysaccharide
MAPK: Mitogen-activated protein kinase
miRNA: MicroRNA
MOI: Multiplicity of infection
MRE: MicroRNA response element
mRNA: Messenger RNA
NFκB: Nuclear factor kappa B
NTD: N-terminal domain
PBS: Phosphate-buffered saline
PD: Periodontal disease
PDL: Periodontal ligament
PFA: Paraformaldehyde
qPCR: Quantitative polymerase chain reaction
Ras: Renin–angiotensin system
RBD: Receptor-binding domain
RIN: RNA integrity number
RISC: RNA-induced silencing complex
RNA: Ribonucleic acid
RT-qPCR: Reverse transcription quantitative polymerase chain reaction
S protein: Spike protein
SARS-CoV-2: Severe acute respiratory syndrome coronavirus 2
SD1-S2: Subdomain 1–subdomain 2
TLR: Toll-like receptor
TLR4: Toll-like receptor 4
TMPRSS2: Transmembrane serine protease 2
TNF: Tumor necrosis factor

## ACKNOWLEDGEMENTS

We thank all our study participants for contributing their clinical data and samples, “Miles Square Health Center” and “University of Illinois Chicago College of Dentistry” clinic staff who supported the recruitment and sample collection. We thank Dr. Lijun Rong and Jazmin Galvan Achi for providing the ACE2-A549 cells. We also thank Dr. Zhiyi Liu at LC Sciences for helping us perform miRNA sequencing experiments.

## DATA AVAILABILITY STATEMENT

The datasets generated and/or analyzed during the current study are available from the corresponding author on reasonable request.

## Notes

### Competing Interest Statement

The authors have declared no competing interest.

