## Supplementary Figure 1 for "Salivary microRNA Profiling of Long COVID Subjects Reveals Host-Encoded Regulators of Inflammation and Viral Persistence"

```
CLUSTAL multiple sequence alignment by MUSCLE (3.8)


BetapUNOSpike         ATGTTTGTGTTCTTGGTGTTGCTTCCACTGGTCAGTTCCCAATGCGTTAATTTTACCACC
OmicronpUNOSpike      ATGTTTGTGTTCTTGGTGTTGCTTCCACTGGTCAGTTCCCAATGCGTTAATCTCACCACC
                      *************************************************** * ******

BetapUNOSpike         CGAACTCAACTCCCACCCGCATATACAAATTCCTTCACCAGAGGAGTGTACTATCCTGAC
OmicronpUNOSpike      CGAACTCAACTCCCACCCGCATATACAAATTCCTTCACCAGAGGAGTGTACTATCCTGAC
                      ************************************************************

BetapUNOSpike         AAAGTGTTTCGGTCAAGTGTCCTCCACTCTACTCAGGACCTCTTTCTGCCTTTCTTTTCT
OmicronpUNOSpike      AAAGTGTTTCGGTCAAGTGTCCTCCACTCTACTCAGGACCTCTTTCTGCCTTTCTTTTCT
                      ************************************************************

BetapUNOSpike         AACGTTACATGGTTTCATGCAATCCATGTGTCTGGGACAAACGGCACCAAACGCTTCGCT
OmicronpUNOSpike      AACGTTACATGGTTTCATGTGATC------TCTGGGACAAACGGCACCAAACGCTTCGAC
                      *******************  ***      ****************************

BetapUNOSpike         AACCCTGTATTGCCATTCAATGATGGGGTGTACTTTGCCTCCACAGAGAAATCCAACATC Spike NTD
OmicronpUNOSpike      AACCCTGTATTGCCATTCAATGATGGGGTGTACTTTGCCTCCATCGAGAAATCCAACATC hsa-miR-106b-5p
                      *******************************************  ***************

BetapUNOSpike         ATTCGAGGATGGATTTTCGGGACTACTCTGGACTCAAAGACACAGAGCCTGCTGATCGTT
OmicronpUNOSpike      ATTCGAGGATGGATTTTCGGGACTACTCTGGACTCAAAGACACAGAGCCTGCTGATCGTT
                      ************************************************************

BetapUNOSpike         AACAACGCCACAAACGTTGTCATCAAAGTGTGCGAATTCCAGTTTTGCAATGATCCCTTC
OmicronpUNOSpike      AACAACGCCACAAACGTTGTCATCAAAGTGTGCGAATTCCAGTTTTGCAATGATCCCTTC
                      ************************************************************

BetapUNOSpike         CTGGGAGTGTACTATCACAAGAATAACAAGTCCTGGATGGAGAGCGAATTTCGGGTCTAC
OmicronpUNOSpike      CTGG---------ACCACAAGAATAACAAGTCCTGGATGGAGAGCGAATTTCGGGTCTAC
                      ****         * *********************************************

BetapUNOSpike         AGCAGCGCAAACAACTGCACCTTCGAGTACGTGAGTCAACCCTTTCTGATGGACCTGGAA
OmicronpUNOSpike      AGCAGCGCAAACAACTGCACCTTCGAGTACGTGAGTCAACCCTTTCTGATGGACCTGGAA
                      ************************************************************

BetapUNOSpike         GGGAAACAGGGAAACTTCAAGAACCTGAGAGAGTTTGTCTTTAAGAACATCGACGGCTAT
OmicronpUNOSpike      GGGAAACAGGGAAACTTCAAGAACCTGAGAGAGTTTGTCTTTAAGAACATCGACGGCTAT
                      ************************************************************

BetapUNOSpike         TTTAAGATCTATAGTAAGCATACGCCTATCAACCTGGTAAG---------GGGTCTTCCC
OmicronpUNOSpike      TTTAAGATCTATAGTAAGCATACGCCTATCA---TTGTAAGGGAGCCCGAGGATCTTCCC
                      *******************************   * *****         ** *******

BetapUNOSpike         CAGGGCTTTTCAGCCCTGGAACCTTTGGTTGACTTGCCTATTGGTATCAATATCACCAGA
OmicronpUNOSpike      CAGGGCTTTTCAGCCCTGGAACCTTTGGTTGACTTGCCTATTGGTATCAATATCACCAGA
                      ************************************************************

BetapUNOSpike         TTTCAGACCCTT---------CATCGGTCTTATCTTACTCCAGGTGATTCCTCCTCCGGG
OmicronpUNOSpike      TTTCAGACCCTTCTGGCATTGCATCGGTCTTATCTTACTCCAGGTGATTCCTCCTCCGGG
                      ************         ***************************************

BetapUNOSpike         TGGACTGCCGGCGCCGCTGCCTACTATGTCGGCTATCTGCAACCAAGAACGTTCCTGCTC
OmicronpUNOSpike      TGGACTGCCGGCGCCGCTGCCTACTATGTCGGCTATCTGCAACCAAGAACGTTCCTGCTC
                      ************************************************************

BetapUNOSpike         AAGTACAACGAAAACGGCACTATTACGGATGCTGTTGATTGTGCCCTGGACCCTCTGTCT
OmicronpUNOSpike      AAGTACAACGAAAACGGCACTATTACGGATGCTGTTGATTGTGCCCTGGACCCTCTGTCT
                      ************************************************************

BetapUNOSpike         GAGACTAAATGCACCCTCAAGAGCTTTACCGTTGAGAAGGGGATTTACCAAACCAGTAAT
```

```
OmicronpUNOSpike      GAGACTAAATGCACCCTCAAGAGCTTTACCGTTGAGAAGGGGATTTACCAAACCAGTAAT
                      ************************************************************

BetapUNOSpike         TTCCGGGTCCAACCCACCGAAAGCATTGTGCGGTTCCCAAATATCACCAATCTGTGTCCC
OmicronpUNOSpike      TTCCGGGTCCAACCCACCGAAAGCATTGTGCGGTTCCCAAATATCACCAATCTGTGTCCC
                      ************************************************************

BetapUNOSpike         TTTGGCGAAGTGTTCAATGCTACAAGGTTTGCTTCTGTGTACGCATGGAATAGGAAACGC
OmicronpUNOSpike      TTTGATGAAGTGTTCAATGCTACAAGGTTTGCTTCTGTGTACGCATGGAATAGGAAACGC
                      ****  ******************************************************

BetapUNOSpike         ATCTCCAATTGTGTCGCTGATTACTCCGTGCTGTACAATTCCGCCTCTTTCTCAACCTTC
OmicronpUNOSpike      ATCTCCAATTGTGTCGCTGATTACTCCGTGCTGTACAATCTGGCCCCATTCTTCACCTTC
                      ***************************************   *** * ****  ******

BetapUNOSpike         AAGTGTTATGGCGTTTCACCTACCAAACTTAACGACCTGTGCTTCACTAATGTGTATGCC
OmicronpUNOSpike      AAGTGTTATGGCGTTTCACCTACCAAACTTAACGACCTGTGCTTCACTAATGTGTATGCC
                      ************************************************************

BetapUNOSpike         GACTCTTTTGTGATACGAGGCGATGAAGTGAGACAGATTGCACCAGGGCAGACCGGCAAT
OmicronpUNOSpike      GACTCTTTTGTGATACGAGGCGATGAAGTGAGACAGATTGCACCAGGGCAGACCGGCAAC
                      ***********************************************************

BetapUNOSpike         ATTGCCGACTACAACTACAAGCTTCCAGATGACTTTACCGGATGTGTTATTGCATGGAAC
OmicronpUNOSpike      ATTGCCGACTACAACTACAAGCTTCCAGATGACTTTACCGGATGTGTTATTGCATGGAAC
                      ************************************************************

BetapUNOSpike         TCAAACAATCTGGATTCCAAGGTGGGTGGCAACTATAACTACCTGTATAGACTGTTCAGG
OmicronpUNOSpike      TCAAACAAGCTGGATTCCAAGGTGAGCGGCAACTATAACTACCTGTATAGACTGTTCAGG
                      ******** *************** * *********************************

BetapUNOSpike         AAATCCAACCTGAAACCATTCGAGCGAGATATAAGCACAGAAATCTACCAGGCTGGAAGT
OmicronpUNOSpike      AAATCCAACCTGAAACCATTCGAGCGAGATATAAGCACAGAAATCTACCAGGCTGGAAAC
                      **********************************************************

BetapUNOSpike         ACGCCCTGCAACGGCGTGAAAGGGTTCAACTGCTACTTCCCATTGCAGAGTTACGGATTC
OmicronpUNOSpike      AAACCCTGCAACGGCGTGGCTGGGTTCAACTGCTACTTCCCATTGCGCAGTTACAGCTTC
                      *  ***************   *************************  ****** * ***

BetapUNOSpike         CAGCCTACATATGGGGTGGGTTACCAACCCTATCGTGTCGTAGTCCTGAGTTTTGAGCTC   hsa-miR-425-5p
OmicronpUNOSpike      AGACCTACATACGGGGTGGGTCACCAACCCTATCGTGTCGTAGTCCTGAGTTTTGAGCTC   hsa-miR-652-3p
                         ******** ********* **************************************   Spike RBD

BetapUNOSpike         CTCCATGCCCCAGCCACAGTCTGTGGCCCCAAGAAAAGCACCAATCTGGTGAAGAACAAA
OmicronpUNOSpike      CTCCATGCCCCAGCCACAGTCTGTGGCCCCAAGAAAAGCACCAATCTGGTGAAGAACAAA
                      ************************************************************

BetapUNOSpike         TGCGTGAACTTTAACTTTAACGGACTCACAGGAACCGGCGTATTGACGGAGAGTAACAAG
OmicronpUNOSpike      TGCGTGAACTTTAACTTTAACGGACTCAAGGGAACCGGCGTATTGACGGAGAGTAACAAG
                      ****************************  ******************************

BetapUNOSpike         AAGTTCCTGCCATTCCAGCAGTTCGGTCGCGATATTGCCGACACTACCGACGCTGTCCGA
OmicronpUNOSpike      AAGTTCCTGCCATTCCAGCAGTTCGGTCGCGATATTGCCGACACTACCGACGCTGTCCGA
                      ************************************************************

BetapUNOSpike         GATCCCCAGACATTGGAGATTCTTGATATCACACCCTGTAGTTTCGGCGGAGTGAGCGTG
OmicronpUNOSpike      GATCCCCAGACATTGGAGATTCTTGATATCACACCCTGTAGTTTCGGCGGAGTGAGCGTG
                      ************************************************************

BetapUNOSpike         ATTACGCCCGGAACCAATACCAGCAATCAGGTTGCCGTCCTGTATCAGGGTGTGAATTGC
OmicronpUNOSpike      ATTACGCCCGGAACCAATACCAGCAATCAGGTTGCCGTCCTGTATCAGGGCGTGAATTGC
                      ************************************************** *********

BetapUNOSpike         ACCGAGGTACCTGTCGCCATCCACGCTGACCAACTTACACCCACATGGCGAGTATATTCC
OmicronpUNOSpike      ACCGAGGTACCTGTCGCCATCCACGCTGACCAACTTACACCCACATGGCGAGTATATTCC
```

```
                      ************************************************************

BetapUNOSpike         ACCGGCTCCAACGTCTTTCAGACACGTGCTGGATGTCTGATCGGTGCAGAACACGTTAAT
OmicronpUNOSpike      ACCGGCTCCAACGTCTTTCAGACACGTGCTGGATGTCTGATCGGTGCAGAATATGTTAAT
                      *************************************************** * ******

BetapUNOSpike         AATAGCTACGAGTGTGATATCCCCATCGGTGCTGGAATATGCGCCTCTTATCAAACTCAA
OmicronpUNOSpike      AATAGCTACGAGTGTGATATCCCCATCGGTGCTGGAATATGCGCCTCTTATCAAACTCAA
                      ************************************************************

BetapUNOSpike         ACCAACTCTCCTAGGCGGGCACGTAGTGTAGCATCCCAAAGTATCATTGCCTACACAATG
OmicronpUNOSpike      ACCAAATCTCACAGGCGGGCACGTAGTGTAGCATCCCAAAGTATCATTGCCTACACAATG
                      ***** ****  ************************************************

BetapUNOSpike         AGCCTCGGTGTAGAGAATTCTGTCGCCTACAGCAACAACTCCATTGCTATCCCTACTAAC
OmicronpUNOSpike      AGCCTCGGTGCTGAGAATTCTGTCGCCTACAGCAACAACTCCATTGCTATCCCTACTAAC
                      **********  ************************************************

BetapUNOSpike         TTCACAATCAGTGTGACAACTGAAATTCTGCCCGTATCTATGACCAAAACAAGCGTTGAC
OmicronpUNOSpike      TTCACAATCAGTGTGACAACTGAAATTCTGCCCGTATCTATGACCAAAACAAGCGTTGAC
                      ************************************************************

BetapUNOSpike         TGCACCATGTACATCTGTGGCGATTCTACCGAATGTAGCAATCTCCTCCTGCAATACGGA
OmicronpUNOSpike      TGCACCATGTACATCTGTGGCGATTCTACCGAATGTAGCAATCTCCTCCTGCAATACGGA
                      ************************************************************

BetapUNOSpike         TCATTCTGCACTCAGCTGAATCGTGCCCTCACAGGTATTGCAGTTGAGCAGGACAAGAAT
OmicronpUNOSpike      TCATTCTGCACTCAGCTGAAGCGTGCCCTCACAGGTATTGCAGTTGAGCAGGACAAGAAT
                      ******************** ***************************************

BetapUNOSpike         ACGCAGGAAGTGTTTGCCCAGGTGAAGCAAATCTACAAAACTCCACCCATAAAAGACTTT
OmicronpUNOSpike      ACGCAGGAAGTGTTTGCCCAGGTGAAGCAAATCTACAAAACTCCACCCATAAAATACTTT
                      ****************************************************** *****

BetapUNOSpike         GGCGGATTCAATTTCTCACAGATCCTGCCCGATCCCTCAAAACCCTCCAAGCGTAGCTTT
OmicronpUNOSpike      GGCGGATTCAATTTCTCACAGATCCTGCCCGATCCCTCAAAACCCTCCAAGCGTAGCTTT
                      ************************************************************

BetapUNOSpike         ATCGAGGATCTGCTCTTCAACAAGGTAACCCTCGCAGATGCCGGTTTCATCAAGCAGTAT
OmicronpUNOSpike      ATCGAGGATCTGCTCTTCAACAAGGTAACCCTCGCAGATGCCGGTTTCATCAAGCAGTAT
                      ************************************************************

BetapUNOSpike         GGCGATTGTCTGGGAGACATCGCCGCTCGGGACCTGATCTGTGCACAGAAGTTCAATGGA
OmicronpUNOSpike      GGCGATTGTCTGGGAGACATCGCCGCTCGGGACCTGATCTGTGCACAGAAGTTCAAAGGA
                      ******************************************************** ***

BetapUNOSpike         CTGACCGTGCTGCCTCCCTTGCTGACCGACGAGATGATAGCCCAATACACTAGCGCCCTG
OmicronpUNOSpike      CTGACCGTGCTGCCTCCCTTGCTGACCGACGAGATGATAGCCCAATACACTAGCGCCCTG
                      ************************************************************

BetapUNOSpike         CTGGCCGGCACCATCACTTCTGGGTGGACATTCGGAGCTGGCGCTGCCCTTCAGATTCCT
OmicronpUNOSpike      CTGGCCGGCACCATCACTTCTGGGTGGACATTCGGAGCTGGCGCTGCCCTTCAGATTCCT
                      ************************************************************

BetapUNOSpike         TTTGCTATGCAGATGGCCTACCGCTTTAACGGCATCGGTGTGACACAAAACGTTCTGTAT
OmicronpUNOSpike      TTTGCTATGCAGATGGCCTACCGCTTTAACGGCATCGGTGTGACACAAAACGTTCTGTAT
                      ************************************************************

BetapUNOSpike         GAAAACCAGAAACTCATCGCCAACCAGTTCAACAGTGCTATCGGTAAGATACAGGATAGC
OmicronpUNOSpike      GAAAACCAGAAACTCATCGCCAACCAGTTCAACAGTGCTATCGGTAAGATACAGGATAGC
                      ************************************************************

BetapUNOSpike         CTGTCATCCACTGCCAGCGCATTGGGAAAGTTGCAGGATGTAGTGAACCAGAATGCCCAG
OmicronpUNOSpike      CTGTCATCCACTGCCAGCGCATTGGGAAAGTTGCAGGATGTAGTGAACCACAATGCCCAG
                      ************************************************** *********
```

```
BetapUNOSpike           GCACTTAACACCCTGGTGAAACAGCTCTCTTCAAATTTTGGTGCCATTTCTAGCGTGCTG    hsa-miR-652-3p
OmicronpUNOSpike        GCACTTAACACCCTGGTGAAACAGCTCTCTTCAAAGTTTGGTGCCATTTCTAGCGTGCTG
                        *********************************** ************************
                                                          SARS-CoV-like_Spike_SD1-2_S1-S2_S2

BetapUNOSpike           AATGACATACTGAGCCGGTTGGACAAGGTGGAGGCTGAAGTGCAGATTGATAGGCTGATA
OmicronpUNOSpike        AATGACATATTTAGCCGGTTGGACAAGGTGGAGGCTGAAGTGCAGATTGATAGGCTGATA
                        ********* * ************************************************

BetapUNOSpike           ACTGGGCGCCTTCAGTCTCTTCAGACCTATGTGACCCAGCAGCTCATCCGCGCTGCTGAA
OmicronpUNOSpike        ACTGGGCGCCTTCAGTCTCTTCAGACCTATGTGACCCAGCAGCTCATCCGCGCTGCTGAA
                        ************************************************************

BetapUNOSpike           ATTCGCGCATCCGCTAACCTGGCAGCAACCAAAATGTCCGAGTGTGTGCTGGGTCAGTCT
OmicronpUNOSpike        ATTCGCGCATCCGCTAACCTGGCAGCAACCAAAATGTCCGAGTGTGTGCTGGGTCAGTCT
                        ************************************************************
                                                          SARS-CoV-like_Spike_SD1-2_S1-S2_S2

BetapUNOSpike           AAGAGAGTGGACTTTTGCGGGAAGGGGTATCACCTGATGTCTTTTCCTCAGTCTGCACCC
OmicronpUNOSpike        AAGAGAGTGGACTTTTGCGGGAAGGGGTATCACCTGATGTCTTTTCCTCAGTCTGCACCC
                        ************************************************************
                                                                      hsa-miR-106b-3p

BetapUNOSpike           CATGGTGTGGTCTTTCTGCACGTGACTTATGTCCCAGCTCAGGAAAAGAACTTCACTACA
OmicronpUNOSpike        CATGGTGTGGTCTTTCTGCACGTGACTTATGTCCCAGCTCAGGAAAAGAACTTCACTACA
                        ************************************************************

BetapUNOSpike           GCCCCAGCCATCTGCCACGATGGGAAAGCCCACTTTCCCAGGGAAGGCGTATTCGTGTCC
OmicronpUNOSpike        GCCCCAGCCATCTGCCACGATGGGAAAGCCCACTTTCCCAGGGAAGGCGTATTCGTGTCC
                        ************************************************************

BetapUNOSpike           AATGGTACTCATTGGTTCGTCACTCAGAGAAATTTCTACGAGCCCCAGATTATAACCACT
OmicronpUNOSpike        AATGGTACTCATTGGTTCGTCACTCAGAGAAATTTCTACGAGCCCCAGATTATAACCACT
                        ************************************************************

BetapUNOSpike           GACAATACATTTGTATCCGGCAATTGTGATGTGGTTATCGGGATTGTGAATAATACTGTT
OmicronpUNOSpike        GACAATACATTTGTATCCGGCAATTGTGATGTGGTTATCGGGATTGTGAATAATACTGTT
                        ************************************************************

BetapUNOSpike           TACGATCCTTTGCAGCCAGAGCTGGACTCCTTCAAGGAGGAGCTTGACAAATATTTTAAG
OmicronpUNOSpike        TACGATCCTTTGCAGCCAGAGCTGGACTCCTTCAAGGAGGAGCTTGACAAATATTTTAAG
                        ************************************************************

BetapUNOSpike           AATCACACATCACCTGACGTCGACCTCGGAGATATTTCAGGAATCAATGCTTCCGTGGTC
OmicronpUNOSpike        AATCACACATCACCTGACGTCGACCTCGGAGATATTTCAGGAATCAATGCTTCCGTGGTC
                        ************************************************************

BetapUNOSpike           AATATTCAGAAGGAGATAGACAGGCTGAATGAGGTTGCCAAGAACCTCAACGAGTCTCTG
OmicronpUNOSpike        AATATTCAGAAGGAGATAGACAGGCTGAATGAGGTTGCCAAGAACCTCAACGAGTCTCTG
                        ************************************************************

BetapUNOSpike           ATCGATCTGCAGGAGTTGGGCAAGTACGAACAGTATATCAAATGGCCATGGTACATTTGG
OmicronpUNOSpike        ATCGATCTGCAGGAGTTGGGCAAGTACGAACAGTATATCAAATGGCCTTGGTACATTTGG
                        *********************************************** ************

BetapUNOSpike           CTTGGGTTCATTGCTGGGCTGATAGCTATCGTCATGGTGACAATTATGTTGTGTTGCATG
OmicronpUNOSpike        CTTGGGTTCATTGCTGGGCTGATAGCTATCGTCATGGTGACAATTATGTTGTGTTGCATG
                        ************************************************************

BetapUNOSpike           ACATCCTGCTGTAGTTGTCTGAAGGGCTGCTGCTCATGCGGCAGCTGTTGCTAA
OmicronpUNOSpike        ACATCCTGCTGTAGTTGTCTGAAGGGCTGCTGCTCATGCGGCAGCTGTTGCTAA
                        ******************************************************
```
