## Supplementary Figure 2 for "Salivary microRNA Profiling of Long COVID Subjects Reveals Host-Encoded Regulators of Inflammation and Viral Persistence"

Version: RNAhybrid 2.2

Command line:/vol/bioapps/bin/RNAhybrid.bin -f 2,8 -b 4 -e -20 -D -n 22 -m 29688 -t /var/bibiserv2/anonymous/rnahybrid/23/21/04/bibiserv2_2025-08-23_210406_VdAkh/rnahybrid_input_target_rna_sequences_.file -v 2 -q /var/bibiserv2/anonymous/rnahybrid/23/21/04/bibiserv2_2025-08-23_210406_VdAkh/rnahybrid_input_mirna_sequences.file

searching

dataset: 1

mde of hsa-miR-652-3p: -47.799999

mde of hsa-miR-106b-3p: -51.400005

mde of hsa-miR-30e-3p: -43.799999

Individual hits

------------------------------------------------------------

dataset: 1

target: PV917192.1

length: 29688

miRNA : hsa-miR-652-3p

length: 21

mfe: -27.7 kcal/mol

p-value: undefined

position 19381

target 5' U	A	C 3'

GCAAUUU GGUGGUGCUGU

UGUUGGG UCACCGCGGUA

miRNA 3' G	A	A 5'

------------------------------------------------------------

dataset: 1

target: PV917192.1

length: 29688

miRNA : hsa-miR-652-3p

length: 21

mfe: -23.9 kcal/mol

p-value: undefined

position 29145

target 5'	U	ACACA	C 3'

GACCU	GGUGCCAU

UUGGG	CCGCGGUA

miRNA 3' GUG	AUCA	A 5'

------------------------------------------------------------

dataset: 1

target: PV917192.1

length: 29688

miRNA : hsa-miR-652-3p

length: 21

mfe: -23.4 kcal/mol

p-value: undefined

position 8263

target 5'	U	G	C C	A 3'

GACU UAG G GCGUCAU

UUGG AUC C CGCGGUA

miRNA 3' GUG	G	A	A 5'

------------------------------------------------------------

dataset: 1

target: PV917192.1

length: 29688

miRNA : hsa-miR-652-3p

length: 21

mfe: -23.3 kcal/mol

p-value: undefined

position 1978

target 5' A	ACAUUACA	C 3'

AUGGCCU	GGUGGUGUUGUU

UGUUGGG	UCACCGCGGUAA

miRNA 3' G	A	5'

------------------------------------------------------------

dataset: 1

target: PV917192.1

length: 29688

miRNA : hsa-miR-106b-3p

length: 22

mfe: -27.4 kcal/mol

p-value: undefined

position 5656

target 5' U U	ACUGGUAAUUA	U 3'

GC AGUGAGUAC	C CAGUGUGG

CG UCGUUCAUG	G GUCACGCC

miRNA 3'	G	U	5'

------------------------------------------------------------

dataset: 1

target: PV917192.1

length: 29688

miRNA : hsa-miR-106b-3p

length: 22

mfe: -24.2 kcal/mol

p-value: undefined

position 7908

target 5' A	UU	U GAUGUUGGUG	A 3'

GGCA AGUG CU	AUAGUGCGG

UCGU UCAU GG	UGUCACGCC

miRNA 3' CG	G	5'

------------------------------------------------------------

dataset: 1

target: PV917192.1

length: 29688

miRNA : hsa-miR-106b-3p

length: 22

mfe: -23.1 kcal/mol

p-value: undefined

position 28066

target 5' U GU	U 3'

GG AGU CUUGUAGUGCG

UC UCA GGGUGUCACGC

miRNA 3' CG GU	U	C 5'

------------------------------------------------------------

dataset: 1

target: PV917192.1

length: 29688

miRNA : hsa-miR-106b-3p

length: 22

mfe: -21.9 kcal/mol

p-value: undefined

position 29551

target 5' A	U	UAAU	U 3'

UAGCAA	CUU	CAGUGUG

GUCGUU	GGG	GUCACGC

miRNA 3' C	CAU	U	C 5'

------------------------------------------------------------

dataset: 1

target: PV917192.1

length: 29688

miRNA : hsa-miR-30e-3p

length: 22

mfe: -21.3 kcal/mol

p-value: undefined

position 15744

target 5'	A	G	C 3'

AAAUGUU GACUGAGA

UUUGUAG CUGACUUU

miRNA 3' CGACA	G	C 5'

------------------------------------------------------------

dataset: 1

target: PV917192.1

length: 29688

miRNA : hsa-miR-30e-3p

length: 22

mfe: -20.9 kcal/mol

p-value: undefined

position 16515

target 5' A U	C	U	A 3'

GC	AACA CUG ACUGAAAG

CG	UUGU GGC UGACUUUC

miRNA 3'	ACAU	A	5'

------------------------------------------------------------
