## Supplementary Figure 3 for "Salivary microRNA Profiling of Long COVID Subjects Reveals Host-Encoded Regulators of Inflammation and Viral Persistence"

Forward primer to amplify B.1.351 and B.1.1.529 spike sequences:

5’-**GCACTCGAG**GGACCTGCAGGGCCTGAAATA-3’ (XhoI site in bold)

Reverse primer to amplify B.1.351 and B.1.1.529 spike sequences:

5’-**ATGCGGCCGC**TTCGATAGAGAAATGTTCTGGC-3’ (NotI site in bold.

Forward primer to amplify B.1.351 and B.1.1.529 nucleotide sequences:

5’-**GCACTCGAG**ATGTCTGATAATGGACCCCAAA-3’ (XhoI site in bold)

Reverse primer to amplify B.1.351 and B.1.1.529 nucleotide sequences

5’-**ATGCGGCCGC**TTAGGCCTGAGTTGAGTCAG-3’ (NotI site in bold)

**pUNO1-SpikeV11**

(7176 bp)

version 21L09v46

GGACCTGCAGGGCCTGAAATAACCTCTGAAAGAGGAACTTGGTTAGGTACCTTCTGAGGCGGAAAGAACCAGCTGTGGAATGTGTGTCAGTTAGGGTGTGGAAAGTCCCCAGGCTCCCCAGCAGGCAGAAGTATGCAAAGCATGCATCTCAATTAGTCAGCAACCAGGTGTGGAAAGTCCCCAGGCTCCCCAGCAGGCAGAAGTATGCAAAGCATGCATCTCAATTAGTCAGCAACCATAGTCCCACTAGTGCTCCGGTGCCCGTCAGTGGGCAGAGCGCACATCGCCCACAGTCCCCGAGAAGTTGGGGGGAGGGGTCGGCAATTGAACGGGTGCCTAGAGAAGGTGGCGCGGGGTAAACTGGGAAAGTGATGTCGTGTACTGGCTCCGCCTTTTTCCCGAGGGTGGGGGAGAACCGTATATAAGTGCAGTAGTCGCCGTGAACGTTCTTTTTCGCAACGGGTTTGCCGCCAGAACACAGCTGAAGCTTCGAGGGGCTCGCATCTCTCCTTCACGCGCCCGCCGCCCTACCTGAGGCCGCCATCCACGCCGGTTGAGTCGCGTTCTGCCGCCTCCCGCCTGTGGTGCCTCCTGAACTGCGTCCGCCGTCTAGGTAAGTTTAAAGCTCAGGTCGAGACCGGGCCTTTGTCCGGCGCTCCCTTGGAGCCTACCTAGACTCAGCCGGCTCTCCACGCTTTGCCTGACCCTGCTTGCTCAACTCTACGTCTTTGTTTCGTTTTCTGTTCTGCGCAGTTACAGATCCAAGCTGTGACCGGCGCCTACCTGAGATCACCGGTCAACATGTTTGTGTTCTTGGTGTTGCTTCCACTGGTCAGTTCCCAATGCGTTAATCTCACCACCCGAACTCAACTCCCACCCGCATATACAAATTCCTTCACCAGAGGAGTGTACTATCCTGACAAAGTGTTTCGGTCAAGTGTCCTCCACTCTACTCAGGACCTCTTTCTGCCTTTCTTTTCTAACGTTACATGGTTTCATGTGATCTCTGGGACAAACGGCACCAAACGCTTCGACAACCCTGTATTGCCATTCAATGATGGGGTGTACTTTGCCTCCATCGAGAAATCCAACATCATTCGAGGATGGATTTTCGGGACTACTCTGGACTCAAAGACACAGAGCCTGCTGATCGTTAACAACGCCACAAACGTTGTCATCAAAGTGTGCGAATTCCAGTTTTGCAATGATCCCTTCCTGGACCACAAGAATAACAAGTCCTGGATGGAGAGCGAATTTCGGGTCTACAGCAGCGCAAACAACTGCACCTTCGAGTACGTGAGTCAACCCTTTCTGATGGACCTGGAAGGGAAACAGGGAAACTTCAAGAACCTGAGAGAGTTTGTCTTTAAGAACATCGACGGCTATTTTAAGATCTATAGTAAGCATACGCCTATCATTGTAAGGGAGCCCGAGGATCTTCCCCAGGGCTTTTCAGCCCTGGAACCTTTGGTTGACTTGCCTATTGGTATCAATATCACCAGATTTCAGACCCTTCTGGCATTGCATCGGTCTTATCTTACTCCAGGTGATTCCTCCTCCGGGTGGACTGCCGGCGCCGCTGCCTACTATGTCGGCTATCTGCAACCAAGAACGTTCCTGCTCAAGTACAACGAAAACGGCACTATTACGGATGCTGTTGATTGTGCCCTGGACCCTCTGTCTGAGACTAAATGCACCCTCAAGAGCTTTACCGTTGAGAAGGGGATTTACCAAACCAGTAATTTCCGGGTCCAACCCACCGAAAGCATTGTGCGGTTCCCAAATATCACCAATCTGTGTCCCTTTGATGAAGTGTTCAATGCTACAAGGTTTGCTTCTGTGTACGCATGGAATAGGAAACGCATCTCCAATTGTGTCGCTGATTACTCCGTGCTGTACAATCTGGCCCCATTCTTCACCTTCAAGTGTTATGGCGTTTCACCTACCAAACTTAACGACCTGTGCTTCACTAATGTGTATGCCGACTCTTTTGTGATACGAGGCGATGAAGTGAGACAGATTGCACCAGGGCAGACCGGCAACATTGCCGACTACAACTACAAGCTTCCAGATGACTTTACCGGATGTGTTATTGCATGGAACTCAAACAAGCTGGATTCCAAGGTGAGCGGCAACTATAACTACCTGTATAGACTGTTCAGGAAATCCAACCTGAAACCATTCGAGCGAGATATAAGCACAGAAATCTACCAGGCTGGAAACAAACCCTGCAACGGCGTGGCTGGGTTCAACTGCTACTTCCCATTGCGCAGTTACAGCTTCAGACCTACATACGGGGTGGGTCACCAACCCTATCGTGTCGTAGTCCTGAGTTTTGAGCTCCTCCATGCCCCAGCCACAGTCTGTGGCCCCAAGAAAAGCACCAATCTGGTGAAGAACAAATGCGTGAACTTTAACTTTAACGGACTCAAGGGAACCGGCGTATTGACGGAGAGTAACAAGAAGTTCCTGCCATTCCAGCAGTTCGGTCGCGATATTGCCGACACTACCGACGCTGTCCGAGATCCCCAGACATTGGAGATTCTTGATATCACACCCTGTAGTTTCGGCGGAGTGAGCGTGATTACGCCCGGAACCAATACCAGCAATCAGGTTGCCGTCCTGTATCAGGGCGTGAATTGCACCGAGGTACCTGTCGCCATCCACGCTGACCAACTTACACCCACATGGCGAGTATATTCCACCGGCTCCAACGTCTTTCAGACACGTGCTGGATGTCTGATCGGTGCAGAATATGTTAATAATAGCTACGAGTGTGATATCCCCATCGGTGCTGGAATATGCGCCTCTTATCAAACTCAAACCAAATCTCACAGGCGGGCACGTAGTGTAGCATCCCAAAGTATCATTGCCTACACAATGAGCCTCGGTGCTGAGAATTCTGTCGCCTACAGCAACAACTCCATTGCTATCCCTACTAACTTCACAATCAGTGTGACAACTGAAATTCTGCCCGTATCTATGACCAAAACAAGCGTTGACTGCACCATGTACATCTGTGGCGATTCTACCGAATGTAGCAATCTCCTCCTGCAATACGGATCATTCTGCACTCAGCTGAAGCGTGCCCTCACAGGTATTGCAGTTGAGCAGGACAAGAATACGCAGGAAGTGTTTGCCCAGGTGAAGCAAATCTACAAAACTCCACCCATAAAATACTTTGGCGGATTCAATTTCTCACAGATCCTGCCCGATCCCTCAAAACCCTCCAAGCGTAGCTTTATCGAGGATCTGCTCTTCAACAAGGTAACCCTCGCAGATGCCGGTTTCATCAAGCAGTATGGCGATTGTCTGGGAGACATCGCCGCTCGGGACCTGATCTGTGCACAGAAGTTCAAAGGACTGACCGTGCTGCCTCCCTTGCTGACCGACGAGATGATAGCCCAATACACTAGCGCCCTGCTGGCCGGCACCATCACTTCTGGGTGGACATTCGGAGCTGGCGCTGCCCTTCAGATTCCTTTTGCTATGCAGATGGCCTACCGCTTTAACGGCATCGGTGTGACACAAAACGTTCTGTATGAAAACCAGAAACTCATCGCCAACCAGTTCAACAGTGCTATCGGTAAGATACAGGATAGCCTGTCATCCACTGCCAGCGCATTGGGAAAGTTGCAGGATGTAGTGAACCACAATGCCCAGGCACTTAACACCCTGGTGAAACAGCTCTCTTCAAAGTTTGGTGCCATTTCTAGCGTGCTGAATGACATATTTAGCCGGTTGGACAAGGTGGAGGCTGAAGTGCAGATTGATAGGCTGATAACTGGGCGCCTTCAGTCTCTTCAGACCTATGTGACCCAGCAGCTCATCCGCGCTGCTGAAATTCGCGCATCCGCTAACCTGGCAGCAACCAAAATGTCCGAGTGTGTGCTGGGTCAGTCTAAGAGAGTGGACTTTTGCGGGAAGGGGTATCACCTGATGTCTTTTCCTCAGTCTGCACCCCATGGTGTGGTCTTTCTGCACGTGACTTATGTCCCAGCTCAGGAAAAGAACTTCACTACAGCCCCAGCCATCTGCCACGATGGGAAAGCCCACTTTCCCAGGGAAGGCGTATTCGTGTCCAATGGTACTCATTGGTTCGTCACTCAGAGAAATTTCTACGAGCCCCAGATTATAACCACTGACAATACATTTGTATCCGGCAATTGTGATGTGGTTATCGGGATTGTGAATAATACTGTTTACGATCCTTTGCAGCCAGAGCTGGACTCCTTCAAGGAGGAGCTTGACAAATATTTTAAGAATCACACATCACCTGACGTCGACCTCGGAGATATTTCAGGAATCAATGCTTCCGTGGTCAATATTCAGAAGGAGATAGACAGGCTGAATGAGGTTGCCAAGAACCTCAACGAGTCTCTGATCGATCTGCAGGAGTTGGGCAAGTACGAACAGTATATCAAATGGCCTTGGTACATTTGGCTTGGGTTCATTGCTGGGCTGATAGCTATCGTCATGGTGACAATTATGTTGTGTTGCATGACATCCTGCTGTAGTTGTCTGAAGGGCTGCTGCTCATGCGGCAGCTGTTGCTAAAGCTAGCTGGCCAGACATGATAAGATACATTGATGAGTTTGGACAAACCACAACTAGAATGCAGTGAAAAAAATGCTTTATTTGTGAAATTTGTGATGCTATTGCTTTATTTGTAACCATTATAAGCTGCAATAAACAAGTTAACAACAACAATTGCATTCATTTTATGTTTCAGGTTCAGGGGGAGGTGTGGGAGGTTTTTTAAAGCAAGTAAAACCTCTACAAATGTGGTATGGAATTCTAAAATACAGCATAGCAAAACTTTAACCTCCAAATCAAGCCTCTACTTGAATCCTTTTCTGAGGGATGAATAAGGCATAGGCATCAGGGGCTGTTGCCAATGTGCATTAGCTGTTTGCAGCCTCACCTTCTTTCATGGAGTTTAAGATATAGTGTATTTTCCCAAGGTTTGAACTAGCTCTTCATTTCTTTATGTTTTAAATGCACTGACCTCCCACATTCCCTTTTTAGTAAAATATTCAGAAATAATTTAAATACATCATTGCAATGAAAATAAATGTTTTTTATTAGGCAGAATCCAGATGCTCAAGGCCCTTCATAATATCCCCCAGTTTAGTAGTTGGACTTAGGGAACAAAGGAACCTTTAATAGAAATTGGACAGCAAGAAAGCGAGCTTCTAGCTTTAGTTCCTGGTGTACTTGAGGGGGATGAGTTCCTCAATGGTGGTTTTGACCAGCTTGCCATTCATCTCAATGAGCACAAAGCAGTCAGGAGCATAGTCAGAGATGAGCTCTCTGCACATGCCACAGGGGCTGACCACCCTGATGGATCTGTCCACCTCATCAGAGTAGGGGTGCCTGACAGCCACAATGGTGTCAAAGTCCTTCTGCCCGTTGCTCACAGCAGACCCAATGGCAATGGCTTCAGCACAGACAGTGACCCTGCCAATGTAGGCCTCAATGTGGACAGCAGAGATGATCTCCCCAGTCTTGGTCCTGATGGCCGCCCCGACATGGTGCTTGTTGTCCTCATAGAGCATGGTGATCTTCTCAGTGGCGACCTCCACCAGCTCCAGATCCTGCTGAGAGATGTTGAAGGTCTTCATGATGGCCCTCCTATAGTGAGTCGTATTATACTATGCCGATATACTATGCCGATGATTAATTGTCAAAACAGCGTGGATGGCGTCTCCAGCTTATCTGACGGTTCACTAAACGAGCTCTGCTTATATAGACCTCCCACCGTACACGCCTACCGCCCATTTGCGTCAATGGGGCGGAGTTGTTACGACATTTTGGAAAGTCCCGTTGATTTACTAGTCAAAACAAACTCCCATTGACGTCAATGGGGTGGAGACTTGGAAATCCCCGTGAGTCAAACCGCTATCCACGCCCATTGATGTACTGCCAAAACCGCATCATCATGGTAATAGCGATGACTAATACGTAGATGTACTGCCAAGTAGGAAAGTCCCATAAGGTCATGTACTGGGCATAATGCCAGGCGGGCCATTTACCGTCATTGACGTCAATAGGGGGCGTACTTGGCATATGATACACTTGATGTACTGCCAAGTGGGCAGTTTACCGTAAATACTCCACCCATTGACGTCAATGGAAAGTCCCTATTGGCGTTACTATGGGAACATACGTCATTATTGACGTCAATGGGCGGGGGTCGTTGGGCGGTCAGCCAGGCGGGCCATTTACCGTAAGTTATGTAACGCCTGCAGGTTAATTAAGAACATGTGAGCAAAAGGCCAGCAAAAGGCCAGGAACCGTAAAAAGGCCGCGTTGCTGGCGTTTTTCCATAGGCTCCGCCCCCCTGACGAGCATCACAAAAATCGACGCTCAAGTCAGAGGTGGCGAAACCCGACAGGACTATAAAGATACCAGGCGTTTCCCCCTGGAAGCTCCCTCGTGCGCTCTCCTGTTCCGACCCTGCCGCTTACCGGATACCTGTCCGCCTTTCTCCCTTCGGGAAGCGTGGCGCTTTCTCATAGCTCACGCTGTAGGTATCTCAGTTCGGTGTAGGTCGTTCGCTCCAAGCTGGGCTGTGTGCACGAACCCCCCGTTCAGCCCGACCGCTGCGCCTTATCCGGTAACTATCGTCTTGAGTCCAACCCGGTAAGACACGACTTATCGCCACTGGCAGCAGCCACTGGTAACAGGATTAGCAGAGCGAGGTATGTAGGCGGTGCTACAGAGTTCTTGAAGTGGTGGCCTAACTACGGCTACACTAGAAGAACAGTATTTGGTATCTGCGCTCTGCTGAAGCCAGTTACCTTCGGAAAAAGAGTTGGTAGCTCTTGATCCGGCAAACAAACCACCGCTGGTAGCGGTGGTTTTTTTGTTTGCAAGCAGCAGATTACGCGCAGAAAAAAAGGATCTCAAGAAGATCCTTTGATCTTTTCTACGGGGTCTGACGCTCAGTGGAACGAAAACTCACGTTAAGGGATTTTGGTCATGGCTAGTTAATTAACATTTAAATCAGCGGCCGCAATAAAATATCTTTATTTTCATTACATCTGTGTGTTGGTTTTTTGTGTGAATCGTAACTAACATACGCTCTCCATCAAAACAAAACGAAACAAAACAAACTAGCAAAATAGGCTGTCCCCAGTGCAAGTGCAGGTGCCAGAACATTTCTCTATCGAA

**pUNO1-SpikeV3**

(7176 bp)

version 21B19v07

GGACCTGCAGGGCCTGAAATAACCTCTGAAAGAGGAACTTGGTTAGGTACCTTCTGAGGCGGAAAGAACCAGCTGTGGAATGTGTGTCAGTTAGGGTGTGGAAAGTCCCCAGGCTCCCCAGCAGGCAGAAGTATGCAAAGCATGCATCTCAATTAGTCAGCAACCAGGTGTGGAAAGTCCCCAGGCTCCCCAGCAGGCAGAAGTATGCAAAGCATGCATCTCAATTAGTCAGCAACCATAGTCCCACTAGTGCTCCGGTGCCCGTCAGTGGGCAGAGCGCACATCGCCCACAGTCCCCGAGAAGTTGGGGGGAGGGGTCGGCAATTGAACGGGTGCCTAGAGAAGGTGGCGCGGGGTAAACTGGGAAAGTGATGTCGTGTACTGGCTCCGCCTTTTTCCCGAGGGTGGGGGAGAACCGTATATAAGTGCAGTAGTCGCCGTGAACGTTCTTTTTCGCAACGGGTTTGCCGCCAGAACACAGCTGAAGCTTCGAGGGGCTCGCATCTCTCCTTCACGCGCCCGCCGCCCTACCTGAGGCCGCCATCCACGCCGGTTGAGTCGCGTTCTGCCGCCTCCCGCCTGTGGTGCCTCCTGAACTGCGTCCGCCGTCTAGGTAAGTTTAAAGCTCAGGTCGAGACCGGGCCTTTGTCCGGCGCTCCCTTGGAGCCTACCTAGACTCAGCCGGCTCTCCACGCTTTGCCTGACCCTGCTTGCTCAACTCTACGTCTTTGTTTCGTTTTCTGTTCTGCGCAGTTACAGATCCAAGCTGTGACCGGCGCCTACCTGAGATCACCGGTCAACATGTTTGTGTTCTTGGTGTTGCTTCCACTGGTCAGTTCCCAATGCGTTAATTTTACCACCCGAACTCAACTCCCACCCGCATATACAAATTCCTTCACCAGAGGAGTGTACTATCCTGACAAAGTGTTTCGGTCAAGTGTCCTCCACTCTACTCAGGACCTCTTTCTGCCTTTCTTTTCTAACGTTACATGGTTTCATGCAATCCATGTGTCTGGGACAAACGGCACCAAACGCTTCGCTAACCCTGTATTGCCATTCAATGATGGGGTGTACTTTGCCTCCACAGAGAAATCCAACATCATTCGAGGATGGATTTTCGGGACTACTCTGGACTCAAAGACACAGAGCCTGCTGATCGTTAACAACGCCACAAACGTTGTCATCAAAGTGTGCGAATTCCAGTTTTGCAATGATCCCTTCCTGGGAGTGTACTATCACAAGAATAACAAGTCCTGGATGGAGAGCGAATTTCGGGTCTACAGCAGCGCAAACAACTGCACCTTCGAGTACGTGAGTCAACCCTTTCTGATGGACCTGGAAGGGAAACAGGGAAACTTCAAGAACCTGAGAGAGTTTGTCTTTAAGAACATCGACGGCTATTTTAAGATCTATAGTAAGCATACGCCTATCAACCTGGTAAGGGGTCTTCCCCAGGGCTTTTCAGCCCTGGAACCTTTGGTTGACTTGCCTATTGGTATCAATATCACCAGATTTCAGACCCTTCATCGGTCTTATCTTACTCCAGGTGATTCCTCCTCCGGGTGGACTGCCGGCGCCGCTGCCTACTATGTCGGCTATCTGCAACCAAGAACGTTCCTGCTCAAGTACAACGAAAACGGCACTATTACGGATGCTGTTGATTGTGCCCTGGACCCTCTGTCTGAGACTAAATGCACCCTCAAGAGCTTTACCGTTGAGAAGGGGATTTACCAAACCAGTAATTTCCGGGTCCAACCCACCGAAAGCATTGTGCGGTTCCCAAATATCACCAATCTGTGTCCCTTTGGCGAAGTGTTCAATGCTACAAGGTTTGCTTCTGTGTACGCATGGAATAGGAAACGCATCTCCAATTGTGTCGCTGATTACTCCGTGCTGTACAATTCCGCCTCTTTCTCAACCTTCAAGTGTTATGGCGTTTCACCTACCAAACTTAACGACCTGTGCTTCACTAATGTGTATGCCGACTCTTTTGTGATACGAGGCGATGAAGTGAGACAGATTGCACCAGGGCAGACCGGCAATATTGCCGACTACAACTACAAGCTTCCAGATGACTTTACCGGATGTGTTATTGCATGGAACTCAAACAATCTGGATTCCAAGGTGGGTGGCAACTATAACTACCTGTATAGACTGTTCAGGAAATCCAACCTGAAACCATTCGAGCGAGATATAAGCACAGAAATCTACCAGGCTGGAAGTACGCCCTGCAACGGCGTGAAAGGGTTCAACTGCTACTTCCCATTGCAGAGTTACGGATTCCAGCCTACATATGGGGTGGGTTACCAACCCTATCGTGTCGTAGTCCTGAGTTTTGAGCTCCTCCATGCCCCAGCCACAGTCTGTGGCCCCAAGAAAAGCACCAATCTGGTGAAGAACAAATGCGTGAACTTTAACTTTAACGGACTCACAGGAACCGGCGTATTGACGGAGAGTAACAAGAAGTTCCTGCCATTCCAGCAGTTCGGTCGCGATATTGCCGACACTACCGACGCTGTCCGAGATCCCCAGACATTGGAGATTCTTGATATCACACCCTGTAGTTTCGGCGGAGTGAGCGTGATTACGCCCGGAACCAATACCAGCAATCAGGTTGCCGTCCTGTATCAGGGTGTGAATTGCACCGAGGTACCTGTCGCCATCCACGCTGACCAACTTACACCCACATGGCGAGTATATTCCACCGGCTCCAACGTCTTTCAGACACGTGCTGGATGTCTGATCGGTGCAGAACACGTTAATAATAGCTACGAGTGTGATATCCCCATCGGTGCTGGAATATGCGCCTCTTATCAAACTCAAACCAACTCTCCTAGGCGGGCACGTAGTGTAGCATCCCAAAGTATCATTGCCTACACAATGAGCCTCGGTGTAGAGAATTCTGTCGCCTACAGCAACAACTCCATTGCTATCCCTACTAACTTCACAATCAGTGTGACAACTGAAATTCTGCCCGTATCTATGACCAAAACAAGCGTTGACTGCACCATGTACATCTGTGGCGATTCTACCGAATGTAGCAATCTCCTCCTGCAATACGGATCATTCTGCACTCAGCTGAATCGTGCCCTCACAGGTATTGCAGTTGAGCAGGACAAGAATACGCAGGAAGTGTTTGCCCAGGTGAAGCAAATCTACAAAACTCCACCCATAAAAGACTTTGGCGGATTCAATTTCTCACAGATCCTGCCCGATCCCTCAAAACCCTCCAAGCGTAGCTTTATCGAGGATCTGCTCTTCAACAAGGTAACCCTCGCAGATGCCGGTTTCATCAAGCAGTATGGCGATTGTCTGGGAGACATCGCCGCTCGGGACCTGATCTGTGCACAGAAGTTCAATGGACTGACCGTGCTGCCTCCCTTGCTGACCGACGAGATGATAGCCCAATACACTAGCGCCCTGCTGGCCGGCACCATCACTTCTGGGTGGACATTCGGAGCTGGCGCTGCCCTTCAGATTCCTTTTGCTATGCAGATGGCCTACCGCTTTAACGGCATCGGTGTGACACAAAACGTTCTGTATGAAAACCAGAAACTCATCGCCAACCAGTTCAACAGTGCTATCGGTAAGATACAGGATAGCCTGTCATCCACTGCCAGCGCATTGGGAAAGTTGCAGGATGTAGTGAACCAGAATGCCCAGGCACTTAACACCCTGGTGAAACAGCTCTCTTCAAATTTTGGTGCCATTTCTAGCGTGCTGAATGACATACTGAGCCGGTTGGACAAGGTGGAGGCTGAAGTGCAGATTGATAGGCTGATAACTGGGCGCCTTCAGTCTCTTCAGACCTATGTGACCCAGCAGCTCATCCGCGCTGCTGAAATTCGCGCATCCGCTAACCTGGCAGCAACCAAAATGTCCGAGTGTGTGCTGGGTCAGTCTAAGAGAGTGGACTTTTGCGGGAAGGGGTATCACCTGATGTCTTTTCCTCAGTCTGCACCCCATGGTGTGGTCTTTCTGCACGTGACTTATGTCCCAGCTCAGGAAAAGAACTTCACTACAGCCCCAGCCATCTGCCACGATGGGAAAGCCCACTTTCCCAGGGAAGGCGTATTCGTGTCCAATGGTACTCATTGGTTCGTCACTCAGAGAAATTTCTACGAGCCCCAGATTATAACCACTGACAATACATTTGTATCCGGCAATTGTGATGTGGTTATCGGGATTGTGAATAATACTGTTTACGATCCTTTGCAGCCAGAGCTGGACTCCTTCAAGGAGGAGCTTGACAAATATTTTAAGAATCACACATCACCTGACGTCGACCTCGGAGATATTTCAGGAATCAATGCTTCCGTGGTCAATATTCAGAAGGAGATAGACAGGCTGAATGAGGTTGCCAAGAACCTCAACGAGTCTCTGATCGATCTGCAGGAGTTGGGCAAGTACGAACAGTATATCAAATGGCCATGGTACATTTGGCTTGGGTTCATTGCTGGGCTGATAGCTATCGTCATGGTGACAATTATGTTGTGTTGCATGACATCCTGCTGTAGTTGTCTGAAGGGCTGCTGCTCATGCGGCAGCTGTTGCTAAAGCTAGCTGGCCAGACATGATAAGATACATTGATGAGTTTGGACAAACCACAACTAGAATGCAGTGAAAAAAATGCTTTATTTGTGAAATTTGTGATGCTATTGCTTTATTTGTAACCATTATAAGCTGCAATAAACAAGTTAACAACAACAATTGCATTCATTTTATGTTTCAGGTTCAGGGGGAGGTGTGGGAGGTTTTTTAAAGCAAGTAAAACCTCTACAAATGTGGTATGGAATTCTAAAATACAGCATAGCAAAACTTTAACCTCCAAATCAAGCCTCTACTTGAATCCTTTTCTGAGGGATGAATAAGGCATAGGCATCAGGGGCTGTTGCCAATGTGCATTAGCTGTTTGCAGCCTCACCTTCTTTCATGGAGTTTAAGATATAGTGTATTTTCCCAAGGTTTGAACTAGCTCTTCATTTCTTTATGTTTTAAATGCACTGACCTCCCACATTCCCTTTTTAGTAAAATATTCAGAAATAATTTAAATACATCATTGCAATGAAAATAAATGTTTTTTATTAGGCAGAATCCAGATGCTCAAGGCCCTTCATAATATCCCCCAGTTTAGTAGTTGGACTTAGGGAACAAAGGAACCTTTAATAGAAATTGGACAGCAAGAAAGCGAGCTTCTAGCTTTAGTTCCTGGTGTACTTGAGGGGGATGAGTTCCTCAATGGTGGTTTTGACCAGCTTGCCATTCATCTCAATGAGCACAAAGCAGTCAGGAGCATAGTCAGAGATGAGCTCTCTGCACATGCCACAGGGGCTGACCACCCTGATGGATCTGTCCACCTCATCAGAGTAGGGGTGCCTGACAGCCACAATGGTGTCAAAGTCCTTCTGCCCGTTGCTCACAGCAGACCCAATGGCAATGGCTTCAGCACAGACAGTGACCCTGCCAATGTAGGCCTCAATGTGGACAGCAGAGATGATCTCCCCAGTCTTGGTCCTGATGGCCGCCCCGACATGGTGCTTGTTGTCCTCATAGAGCATGGTGATCTTCTCAGTGGCGACCTCCACCAGCTCCAGATCCTGCTGAGAGATGTTGAAGGTCTTCATGATGGCCCTCCTATAGTGAGTCGTATTATACTATGCCGATATACTATGCCGATGATTAATTGTCAAAACAGCGTGGATGGCGTCTCCAGCTTATCTGACGGTTCACTAAACGAGCTCTGCTTATATAGACCTCCCACCGTACACGCCTACCGCCCATTTGCGTCAATGGGGCGGAGTTGTTACGACATTTTGGAAAGTCCCGTTGATTTACTAGTCAAAACAAACTCCCATTGACGTCAATGGGGTGGAGACTTGGAAATCCCCGTGAGTCAAACCGCTATCCACGCCCATTGATGTACTGCCAAAACCGCATCATCATGGTAATAGCGATGACTAATACGTAGATGTACTGCCAAGTAGGAAAGTCCCATAAGGTCATGTACTGGGCATAATGCCAGGCGGGCCATTTACCGTCATTGACGTCAATAGGGGGCGTACTTGGCATATGATACACTTGATGTACTGCCAAGTGGGCAGTTTACCGTAAATACTCCACCCATTGACGTCAATGGAAAGTCCCTATTGGCGTTACTATGGGAACATACGTCATTATTGACGTCAATGGGCGGGGGTCGTTGGGCGGTCAGCCAGGCGGGCCATTTACCGTAAGTTATGTAACGCCTGCAGGTTAATTAAGAACATGTGAGCAAAAGGCCAGCAAAAGGCCAGGAACCGTAAAAAGGCCGCGTTGCTGGCGTTTTTCCATAGGCTCCGCCCCCCTGACGAGCATCACAAAAATCGACGCTCAAGTCAGAGGTGGCGAAACCCGACAGGACTATAAAGATACCAGGCGTTTCCCCCTGGAAGCTCCCTCGTGCGCTCTCCTGTTCCGACCCTGCCGCTTACCGGATACCTGTCCGCCTTTCTCCCTTCGGGAAGCGTGGCGCTTTCTCATAGCTCACGCTGTAGGTATCTCAGTTCGGTGTAGGTCGTTCGCTCCAAGCTGGGCTGTGTGCACGAACCCCCCGTTCAGCCCGACCGCTGCGCCTTATCCGGTAACTATCGTCTTGAGTCCAACCCGGTAAGACACGACTTATCGCCACTGGCAGCAGCCACTGGTAACAGGATTAGCAGAGCGAGGTATGTAGGCGGTGCTACAGAGTTCTTGAAGTGGTGGCCTAACTACGGCTACACTAGAAGAACAGTATTTGGTATCTGCGCTCTGCTGAAGCCAGTTACCTTCGGAAAAAGAGTTGGTAGCTCTTGATCCGGCAAACAAACCACCGCTGGTAGCGGTGGTTTTTTTGTTTGCAAGCAGCAGATTACGCGCAGAAAAAAAGGATCTCAAGAAGATCCTTTGATCTTTTCTACGGGGTCTGACGCTCAGTGGAACGAAAACTCACGTTAAGGGATTTTGGTCATGGCTAGTTAATTAACATTTAAATCAGCGGCCGCAATAAAATATCTTTATTTTCATTACATCTGTGTGTTGGTTTTTTGTGTGAATCGTAACTAACATACGCTCTCCATCAAAACAAAACGAAACAAAACAAACTAGCAAAATAGGCTGTCCCCAGTGCAAGTGCAGGTGCCAGAACATTTCTCTATCGAA
